# The fire ant social chromosome exerts a major influence on genome regulation

**DOI:** 10.1101/2025.02.17.638763

**Authors:** Beryl M. Jones, Alex H. Waugh, Michael A. Catto, Sasha Kay, Karl M. Glastad, Michael A.D. Goodisman, Sarah D. Kocher, Brendan G. Hunt

## Abstract

Supergenes underlying complex trait polymorphisms ensure sets of coadapted alleles remain genetically linked. Despite their prevalence in nature, the mechanisms of supergene effects on genome regulation are poorly understood. In the fire ant *Solenopsis invicta*, a supergene containing over 500 individual genes influences trait variation in multiple castes to collectively underpin a colony level social polymorphism. Here, we present results of an integrative investigation of supergene effects on gene regulation. We present analyses of ATAC-seq data to investigate variation in chromatin accessibility by supergene genotype and STARR-seq data to characterize enhancer activity by supergene haplotype. Integration with gene coexpression analyses, newly mapped intact TEs, and previously identified CNVs, collectively reveal widespread effects of the supergene on chromatin structure, gene transcription, and regulatory element activity, with a genome-wide bias for open chromatin and increased expression in the presence of the derived supergene haplotype, particularly in regions that harbor intact TEs. Integrated consideration of CNVs and regulatory element divergence suggests each evolved in concert to shape the expression of supergene encoded factors, including several transcription factors that may directly contribute to the *trans*-regulatory footprint of a heteromorphic social chromosome. Overall, we show how genome structure in the form of a supergene has wide-reaching effects on gene regulation and gene expression.

## Introduction

Genome structure can have fundamentally important effects on evolution and adaptation. For example, chromosomal inversions can facilitate the evolution of complex trait polymorphisms such as ecotypes (Todesco et al. 2020; Hager et al. 2022; Kay et al. 2022; Matschiner et al. 2022), mating morphs (Mank 2023), and sexes (Branco et al. 2018) by ensuring sets of coadapted alleles remain genetically linked (Dobzhansky and Epling 1948) in what is known as a supergene (Schwander et al. 2014; Thompson and Jiggins 2014). Supergenes have been documented across diverse eukaryotic taxa (Wellenreuther and Bernatchez 2018) and chromosomal inversion polymorphisms are pervasive in natural populations (Dobzhansky and Sturtevant 1938; Huang et al. 2014; Todesco et al. 2020; Harringmeyer and Hoekstra 2022). Despite this prevalence, how supergenes influence genome function beyond limiting recombination and affecting selection on linked regions is not well understood.

Among ants, five independent origins of supergenes are known to be associated with polymorphic life histories and social forms (Kay et al. 2022; Chapuisat 2023; Lajmi et al. 2024). In the red imported fire ant, *Solenopsis invicta*, distinct supergene genotypes control whether a colony contains one egg-laying queen (monogyne social form) or multiple egg-laying queens (polygyne social form) (Ross and Keller 1998; Wang et al. 2013; Yan et al. 2020). This variation in colony social organization arises in conjunction with pronounced supergene effects on caste determination (Buechel et al. 2014), reproductive maturation (DeHeer 2002; Nipitwattanaphon et al. 2013; Waugh et al. 2024), dispersal (DeHeer et al. 1999; Goodisman et al. 2000), male survival (Hettesheimer et al. 2025), and queen acceptance by workers (Keller and Ross 1998; Ross and Keller 2002; Zeng et al. 2022). Monogyne queens invariably are homozygous for the non-inverted *Social B* (*SB*) supergene haplotype, and their colonies reproduce when new queens independently disperse (Ross and Keller 1998; DeHeer et al. 1999; DeHeer 2002; Wang et al. 2013; Yan et al. 2020). Polygyne reproductive queens are heterozygous for *SB* and the inversion containing *Sb* supergene alleles, and their colonies reproduce by budding, with workers always present to support the queen (Vargo and Porter 1989; Ross and Keller 1998; DeHeer et al. 1999; DeHeer 2002; Wang et al. 2013; Yan et al. 2020).

The *Sb* supergene haplotype in fire ants is formed by three large chromosomal inversions and a linked centromeric region of suppressed recombination (Yan et al. 2020). *Sb* spread by introgression to several Solenopsis species within the last million years (Helleu et al. 2022; Stolle et al. 2022), where it underpins polygynous social organization (Yan et al. 2020). *Sb* contains over 500 individual protein coding genes (Wang et al. 2013; Yan et al. 2020) and prior research has revealed genetic differentiation of the *SB* and *Sb* supergene variants, including the presence of non-synonymous substitutions (Pracana et al. 2017; Cohanim et al. 2018; Martinez-Ruiz et al. 2020). Since its origin, *Sb* has also increased in length by over 30% (Stolle et al. 2019), an expansion that coincides with extensive social chromosome linked copy number variation in transposable elements (TEs) and host genes (Pracana et al. 2017; Cohanim et al. 2018; Fontana et al. 2020). Whether and how these TEs and gene copies lead to the striking phenotypic differences between social forms of the fire ant is still largely unknown.

We used a combination of genomic approaches to investigate supergene effects on gene regulation in *S. invicta*. We generated ATAC-seq data from individual worker brains to profile chromatin accessibility, leveraged multiple existing RNA-seq datasets (Chandra et al. 2018; Arsenault et al. 2020; Arsenault et al. 2023) to identify gene expression modules linked to supergene genotype, mapped intact TEs and known copy number variants (CNVs; (Fontana et al. 2020)) to long read genome assemblies (Yan et al. 2020; Helleu et al. 2022), and generated STARR-seq data to characterize enhancer activity of *SB* and *Sb* DNA fragments. Our integrative analyses reveal widespread supergene effects on chromatin remodeling, gene transcription, and regulatory element activity that are heavily biased toward *Sb* activation and often coincide with genomic regions harboring intact TEs. Our results further suggest structural variants and regulatory elements evolved in concert to shape the expression of supergene encoded factors, including several transcription factors (TF) we identify as potential drivers of the *trans*-regulatory footprint of the social chromosome.

## Results and Discussion

### The supergene affects global patterns of transcription

To provide insight into the gene regulatory effects of the polygyne-affiliated *Sb* variant of the fire ant supergene on the social chromosome (chromosome 16; Fig. 1a), we first reanalyzed published RNA-seq data (Table S1). Pairwise comparisons of *SB/Sb* and *SB/SB* gene expression data from worker brains, worker gasters, alate gyne (queen) brains, and queen ovaries (n = 6-8 biological replicates per group; Table S1) illustrate a striking bias among differentially expressed genes (DEGs) toward upregulation in *SB/Sb* relative to *SB/SB* samples (Fig. 1b; Table S2; (Arsenault et al. 2023); *Sb/Sb* individuals are low fitness and were excluded from this analysis (Hallar et al. 2007)). A minority of observed DEGs mapped within the supergene region or exhibited copy number differences between *Sb* and *SB* variants of the social chromosome (Fig. 1c). This pattern is consistent with substantial *trans*-regulatory effects of the supergene (Arsenault et al. 2020).

**Figure 1.**
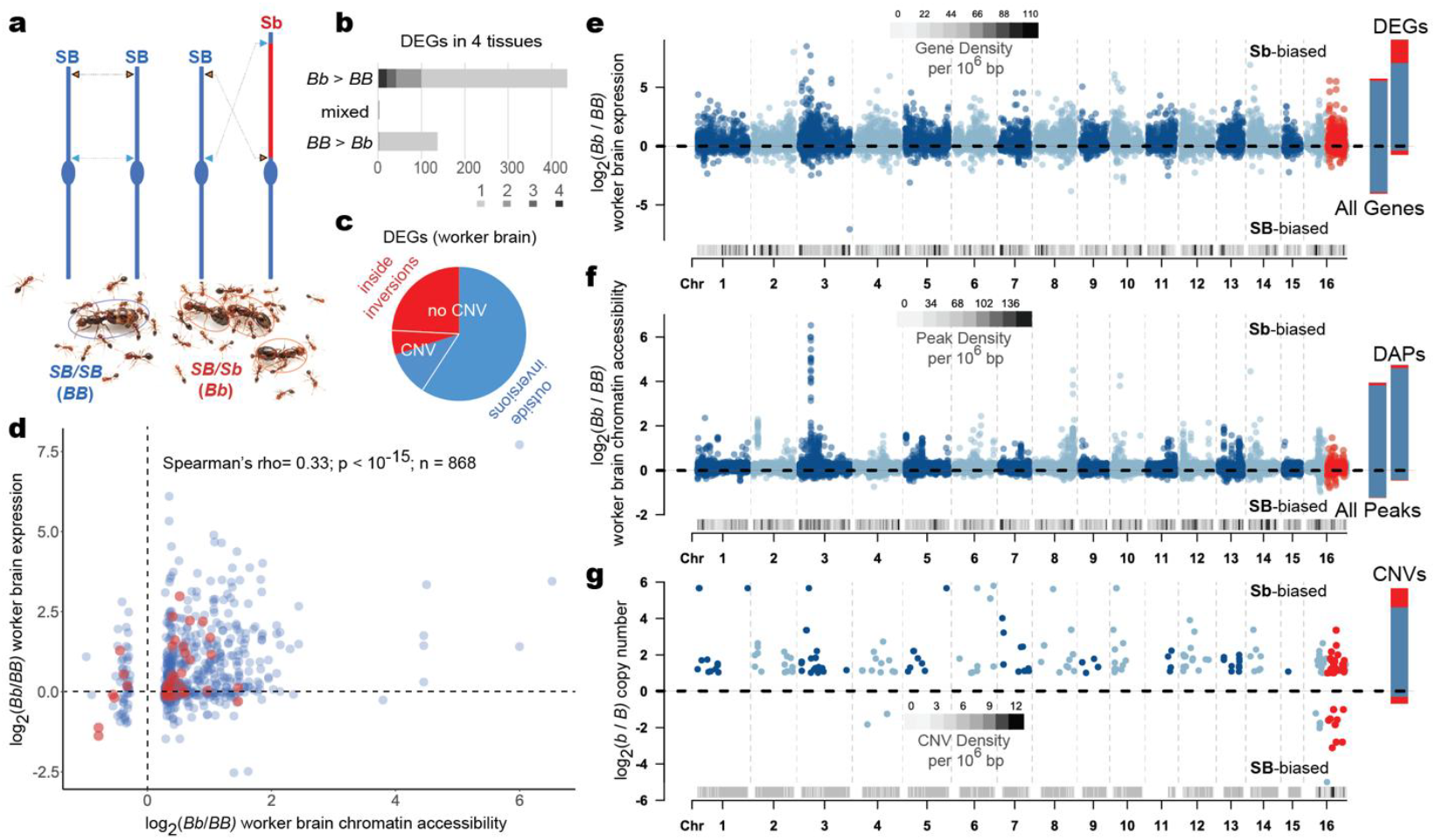
Gene regulatory and copy number variation attributable to the fire ant social chromosome. **(a)** The derived structural variant of chromosome 16 (*Sb*; inversions shown as red chromosomal region; (Helleu et al. 2022)) is restricted to the polygyne social form (having multiple reproductive queens), as opposed to the monogyne social form, of *S. invicta*. Polygyne reproductive queens are heterozygotes for *Sb* and the ancestral structural variant (*SB*), while their offspring are a mix of genotypes (*Sb* homozygotes are scarce). **(b)** Counts of differentially expressed genes (DEGs) according to RNA-seq between *SB/SB* (*BB*) and *SB/Sb* (*Bb*) in worker brains and gasters and alate gyne (queen) brains and ovaries show bias toward upregulation in *SB/Sb* (n = 6-8 replicates per pairwise comparison). **(c)** Proportion of DEGs in worker brains mapped inside or outside the supergene inversion region and with or without *Sb* versus *SB* increased copy number variants (CNVs) shows most DEGs map outside the supergene inversions and lack supergene CNVs. **(d)** Scatterplot of fold difference by supergene genotype in significantly differentially accessible peaks according to ATAC-seq and corresponding RNA-seq fold difference in expression, each from worker brains, shows a significant positive correlation between gene expression and chromatin accessibility with a skew towards increased accessibility and expression in *SB/Sb* relative to *SB/SB*. Genes mapped to the *SB* homologous region of the *Sb* supergene inversions are shown in red. **(e)** log2(*SB/Sb* / *SB/SB*) gene expression level by chromosome position. **(f)** log2(*SB/Sb* / *SB/SB*) chromatin accessibility peaks by chromosome position. **(g)** log2(*Sb / SB*) copy number for CNVs (Fontana et al. 2020) by chromosome position mapped to *SB*. For panels e-g, red dots map to the supergene inversions, greyscale tracks along x-axes show feature density by position, and dashed lines represent a fold-change value of zero.

To assess variation in genome-wide patterns of chromatin accessibility by supergene genotype, we generated ATAC-seq (Buenrostro et al. 2013; Buenrostro et al. 2015) data from individual *SB/Sb* and *SB/SB* worker brains (n = 5 biological replicates, Table S3). Our ATAC-seq data revealed the increase in gene activity in *SB/Sb* (relative to *SB/SB*) is reflected by changes in chromatin accessibility. We observed over 10-fold more significant differentially accessible peaks (DAPs) with higher accessibility in *SB/Sb* than in *SB/SB* (Table S4). We found that DAPs were enriched for motifs of the transcription factors (TFs) DNA replication-related element factor (Dref), Boundary element-associated factory of 32kD (BEAF-32), Erect wing (Ewg), and Pannier (Pnr) with Benjamini-corrected q-value<0.1 (Table S5). These TFs are not located within the supergene, suggesting the enrichment of these TF motifs in DAPs is a downstream consequence of *trans*-regulating factors within the inversion (note that the ortholog of BEAF-32 is unknown in *S. invicta*, but the other TFs are annotated and not within the inverted region). Dref and BEAF-32 are known to physically interact in Drosophila and play a role in chromatin organization, while both Ewg and Pnr are involved in nervous system development and synaptic growth, among other functions (Jenkins et al. 2022). Gene Ontology (GO) annotations of genes near DAPs (937 genes assigned to DAPs based on proximity, see Methods) were enriched for three Biological Processes and two Molecular Functions: “*fatty acid biosynthetic process*”, “*monocarboxylic acid biosynthetic process*”, “*DNA integration*”, “*3-oxoacyl-[acyl-carrier-protein] synthase activity*”, and “*fatty acid synthase activity*”, respectively (FDR<0.1, Table S6). Variation in the regulation of genes that function in fatty acid biosynthesis could potentially contribute to known differences in fat accumulation by gynes (Keller and Ross 1993; Waugh et al. 2024) and variation in the cuticular hydrocarbon profiles of mature queens (Keller and Ross 1998; Zeng et al. 2022) by supergene genotype in *S. invicta*. In this manner, genes involved in fatty acid biosynthesis represent intriguing candidates for pleiotropic effects on trait variation relevant to both nutrient reserve formation and chemical communication.

We observed a highly significant positive correlation between genotypic effects on chromatin accessibility and gene expression levels in worker brains based on an assessment of the 868 DAPs proximal to genes with expression data in the RNA-seq meta-analysis (Fig. 1d). Global visualizations of RNA-seq (Fig. 1e) and ATAC-seq (Fig. 1f) fold-change values between *SB/Sb* and *SB/SB* illustrate a genome-wide distribution of loci with increased gene activity and chromatin accessibility in *SB/Sb* samples. One potentially important contributor to these *trans*-regulatory effects of the relatively young *Sb* structural variant is the existence of many genes throughout the genome with increased copy number in the *Sb* structural variant, apparently via transposable element mediated translocation into and/or duplication within the supergene region of *Sb* (Stolle et al. 2019; Fontana et al. 2020) (CNVs; Fig. 1g; Table S7).

### Constitutive supergene effects are reflected by a small co-expression module with genes from multiple chromosomes

To further characterize the loci differentially regulated by social chromosome genotype, we performed a weighted gene co-expression analysis (Langfelder and Horvath 2008). We included information from 122 RNA-seq libraries (Table S1) including *SB/SB, SB/Sb*, and *Sb/Sb* samples of individual queen and worker tissues from both monogyne and polygyne colonies (Chandra et al. 2018; Arsenault et al. 2020; Arsenault et al. 2023). This resulted in the assignment of 26 gene co-expression modules (Table S8), each of which we examined for enrichment of social chromosome genotype-affiliated differential expression, chromatin accessibility, and GO enrichment (Fig. 2a; Tables S9-S10).

**Figure 2.**
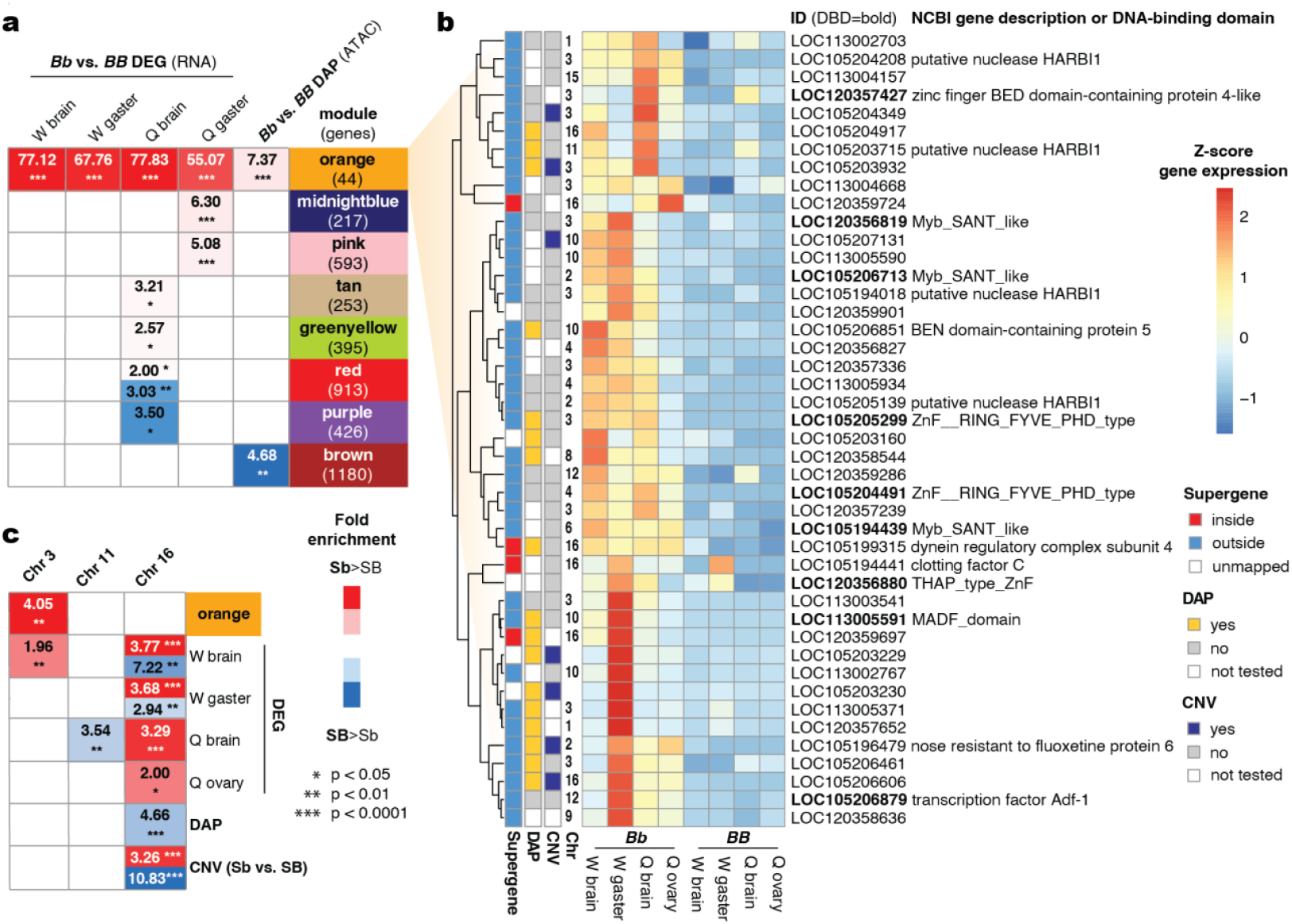
Gene co-expression modules and chromosomes enriched for differential activity by social chromosome genotype. **(a)** WGCNA modules with significant enrichment of DEGs or genes with proximal DAPs. Fold enrichment values are shown relative to expected frequency based on module size, with significance indicated for a hypergeometric overlap test. W, worker; Q, alate gyne (queen). **(b)** Heatmap of standardized expression for genes belonging to the orange gene co-expression module with gene descriptions or included DNA-binding domains (DBD). **(c)** Chromosomes with significant enrichment of DEGs, genes with proximal DAPs, or CNVs by supergene genotype. Fold enrichment values are shown with significance relative to expected frequency based on chromosome gene number indicated for a hypergeometric overlap test. All fold enrichment and p-values are provided in Tables S9 and S11.

The smallest gene co-expression module assigned, orange, contains only 44 genes but is highly enriched for DEGs in all 4 of our focal pairwise RNA-seq comparisons between *SB/Sb* and *SB/SB* (Fig. 2a), containing 19 of 20 4-way overlapping DEGs. Thirty-three of 44 genes from orange are significant DEGs in worker brain, with all 33 of those exhibiting higher expression in *SB/Sb* versus *SB/SB* samples. Orange is also significantly enriched for DAPs between *SB/Sb* and *SB/SB* worker brains (Fig. 2a). Only 7 of the 36 orange module genes with CNV data (Fontana et al. 2020) exhibit supergene-linked copy number variation (Fig. 2b), highlighting that a minority of loci with constitutive expression responsiveness to *Sb* are supergene-linked CNVs. Thus, constitutive expression effects of *Sb* are usually not the result of a simple dosage effect of supergene-linked copy number variation (Fontana et al. 2020). Instead, our results suggest constitutive changes in expression may most commonly be caused by *trans*-regulatory links between supergene encoded factors and genes residing on other chromosomes. Among the 44 genes in orange, 6 are mapped to chromosome 16, the supergene-containing social chromosome, 5 are located on unplaced scaffolds that potentially belong to chromosome 16, and 13 are mapped to chromosome 3, which is devoid of polygyne-affiliated structural variants (Yan et al. 2020) (Fig. 2b). Chromosome 3, like chromosome 16, is significantly enriched for worker brain DEGs. Moreover, chromosome 3 is the only chromosome exhibiting significant enrichment of genes belonging to the orange module following multiple test correction (Fig. 2c, Table S11). The fact that the constitutively *Sb*-responsive orange co-expression module is not enriched for genes mapped to the social chromosome but is enriched for genes mapped to a different chromosome illustrates that the *Sb* supergene haplotype does exhibit strong *trans*-regulatory links to the remainder of the genome.

### Genomic regions with high intact TE density may be hotspots of social chromosome regulatory effects

Transposons can give rise to new genes through “molecular domestication” (Volff 2006). The orange coexpression module includes four distinct (non-CNV) copies of genes encoding the transposon derived protein HARBI1, none of which map to chromosome 16 (Fig. 2b). In a vertebrate study, HARBI1 and a Myb-like protein (including a SANT/Myb/trihelix domain that binds to DNA) were both shown to be derived from a single *Harbinger* transposase (Sinzelle et al. 2008). Remarkably, three distinct (non-CNV) uncharacterized proteins with Myb_SANT_like DNA binding domains (Pfam PF13837 or InterPro IPR028002), none of which map to chromosome 16, are present among the 44 orange module genes. This suggests the expression of multiple HARBI1 and Myb-like proteins are subject to coordinated activation by *Sb* encoded factors. The SANT/Myb/trihelix motif has also been found to function as a DNA binding domain for many transcription factors (Kapitonov and Jurka 2004), which may play a role in this activation.

Based on the domesticated transposons belonging to the supergene responsive orange coexpression module and the importance of transposable elements (TEs) to the formation (Yan et al. 2020) and size expansion (Stolle et al. 2019; Fontana et al. 2020) of *Sb*, we were interested in investigating whether variation in the TE landscape intersected with regional variation in the gene regulatory landscape by supergene genotype. Transcription factor binding sites are frequently embedded within TEs (Sundaram et al. 2014) and TEs may be a frequent source of gene regulatory network innovations (Feschotte 2008; Chuong et al. 2017). Thus, we mapped structurally intact TEs (Ou et al. 2019) in the genome of *S. invicta* to assess whether proximity to TEs is related to whether chromatin exhibits differential accessibility and genes exhibit differentially expression by supergene genotype. We observed that intact TE density is positively correlated with supergene-differentiated DAPs, CNVs, and DEGs across the genome. This relationship is strongest for DAPs, and in each case is strongest when considering large, 1 megabase genomic windows (Table 1).

**Table 1.** Kendall’s rank correlation coefficients (tau) between intact TE counts and supergene associated regulation or CNVs, with partial correlations controlling for gene counts.

| Variable (X) | Correlation tau $X_i$ TE count | Partial correlation tau $X_i$ TE count \| gene count |
| --- | --- | --- |
| **1 Mb windows** |  |  |
| DEG count <sup>a</sup> | 0.415** | 0.266** |
| DAP count <sup>b</sup> | 0.572** | 0.483** |
| CNV count <sup>c</sup> | 0.387** | 0.329** |
| **100 Kb windows** |  |  |
| DEG count | 0.027 | 0.020 |
| DAP count | 0.160** | 0.157** |
| CNV count | 0.108* | 0.106** |

\* $p < 10^{-10}$ ; \*\* $p < 10^{-15}$
<sup>a</sup> Counted as DEG if 1 or more of 4 focal comparisons (Fig. 1b) is significant.
<sup>b</sup> All ATAC-seq DAPs between *Bb* and *BB* worker brains are reported, regardless of proximity to genes.
<sup>c</sup> CNVs from (Fontana et al. 2020) mapped to long read genome builds.

In line with the positive correlation between TE density and DAPs, a visual examination of chromosome 3 (enriched for orange module genes) illustrates an apparent hotspot of intact TEs in a region with many DEGs, DAPs, and CNVs (Fig. 3a). The centromere-proximal region of the social chromosome also exhibits moderate to high intact TE density and harbors many DEGs, DAPs, and CNVs (Fig. 3b, left of red lines). By contrast, the region of the *SB* supergene haplotype that is homologous to the *Sb* chromosomal inversion region coincides with a paucity of TEs (Fig. 3b, between red lines). We used a reference genome that includes the *SB* haplotype of the social chromosome for most of our analyses because the currently available *Sb* genome assembly is less complete and does not have equivalent gene annotations (however, we do consider effects of alternative genome assemblies on our results below). Following its origin, the *Sb* supergene haplotype did accumulate many intact TEs (Stolle et al. 2019; Yan et al. 2020), as confirmed by a comparison of methodologically equivalent long read genome assemblies for *SB* and *Sb* social chromosome haplotypes (which we refer to as v2 (Yan et al. 2020); Fig. 3b-c). Prior optical mapping and kmer analyses suggest the full extent of *Sb* expansion represents an increase in length of over 30% relative to *SB* (Stolle et al. 2019). Overall, these results provide preliminary evidence that TE density may be important to the patterns of differential gene expression and chromatin accessibility observed by supergene genotype, including but not limited to TEs within the expanded *Sb* supergene haplotype.

**Figure 3.**
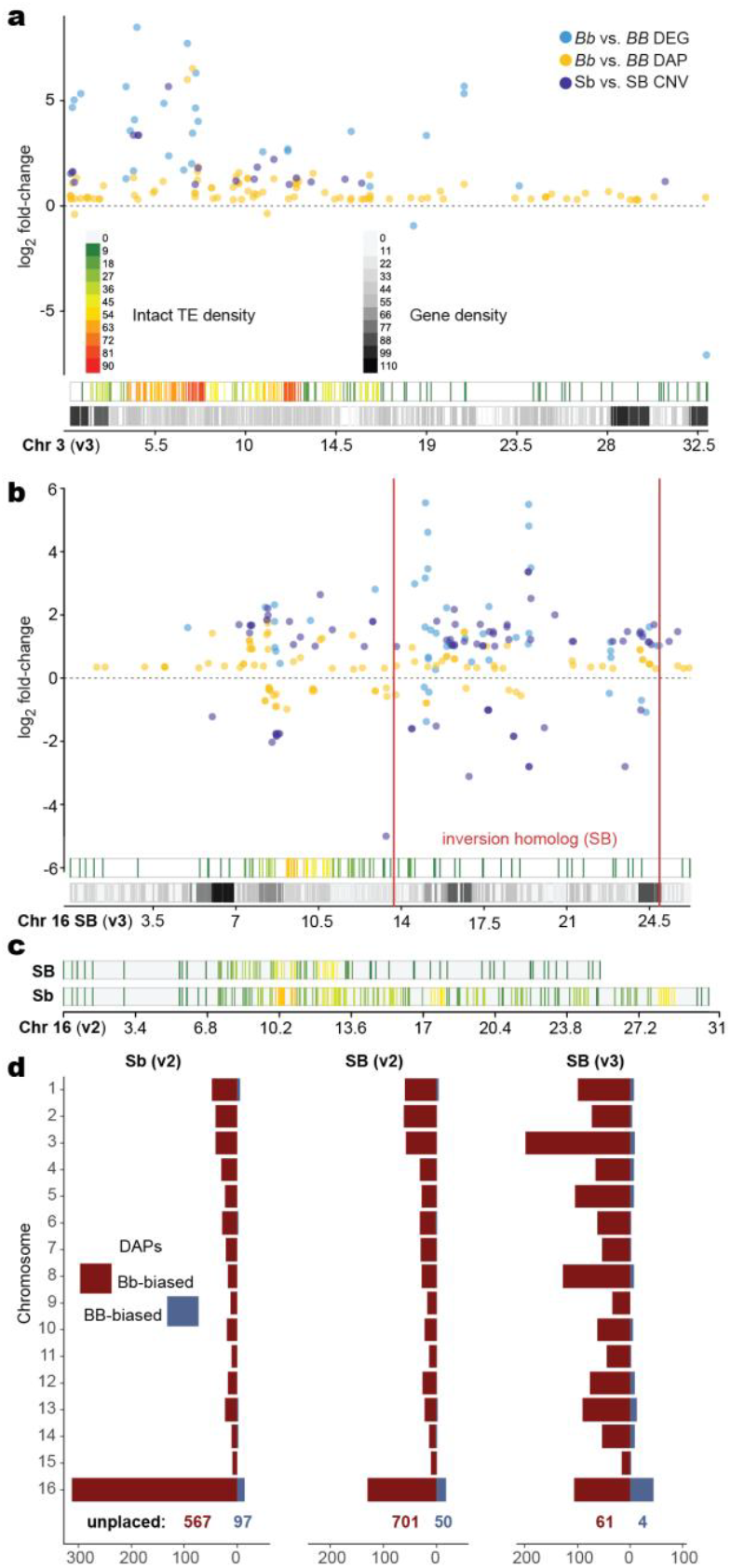
Intact TE density and the genomic distribution of social chromosome regulatory effects. **(a)** Chromosome (Chr) 3 location of DEGs, DAPs, and CNVs with respect to genes and intact TEs. **(b)** *SB* Chr 16 location of DEGs, DAPs, and CNVs with respect to genes and intact TEs. Red lines demark the external bounds of supergene inversion breakpoints in *Sb*. **(c)** Intact TE density for *SB* and *Sb* long read social chromosome assemblies illustrates rapid evolutionary gain of TEs in *Sb* relative to *SB*. **(d)** Distribution of DAPs by chromosome on comparable *Sb* and *SB* long read genome assemblies (v2; (Yan et al. 2020)) and an improved *SB* assembly (v3; (Helleu et al. 2022)) show *trans*-regulatory effects of the supergene and prevalence of DAPs on the social chromosome (Chr 16) and unplaced scaffolds.

Across the genome, genes with evidence of *Sb*-linked CNVs (Fontana et al. 2020) also exhibited a significantly increased overlap with intact TEs as compared to non-CNVs (Fig. S1), with 19.47% of CNVs overlapping at least one intact TE as compared to 6.41% of non-CNVs overlapping at least one intact TE (Fisher’s Exact test, odds ratio=3.6, p=1.3e-12). This suggests TEs are likely to have played a direct role in the translocation and duplication of host genes in the social chromosome, consistent with the importance of TEs to the origins of CNVs more generally (Cerbin and Jiang 2018) and observations of TEs flanking some of the CNVs (Fontana et al. 2020) and one of the large inversions in the fire ant supergene (Yan et al. 2020).

Structural differentiation of heteromorphic chromosomes complicates analyses of gene regulation because cells of diploid organisms include two chromosome variants whereas most analysis pipelines involve mapping to a single reference genome to identify differential activity. To consider such effects on our analyses and interpretation, we assessed how the chromosomal distribution of differentially accessible chromatin peaks by supergene genotype might be influenced by mapping our ATAC-seq data to an *Sb* versus *SB* reference genome assembly and *SB* reference assemblies of variable completeness. Mapping to an *Sb* reference assembly resulted in over twice as many DAPs being assigned to chromosome 16 as compared to the comparable *SB* assembly (Fig. 3d; Fig. S2; Table S12). This suggests some DAPs that were on unplaced scaffolds or mapped to other chromosomes in an *SB* reference genome may actually be *Sb*-linked CNVs with new duplicates present in the supergene (at present, we do not have the chromosome level assembly and gene models for *Sb* needed to fully disentangle this issue directly). When we analyze DAPs mapped to the *Sb* (v2) genome, which are not dependent on gene annotations, the signal of the hotspot of intact TEs and DAPs in chromosome 3 (Fig. 3a) remain intact (Fig. S2-S3). Moreover, the most recent assembly of *SB* shows great reduction in unmapped content (v3, available for only *SB* (Helleu et al. 2022)) as well as increases in *Sb*-biased DAPs in many chromosomes other than the social chromosome (Fig. 1f, Fig. 3d) relative to the less complete *SB* assembly (v2 (Yan et al. 2020)), consistent with our hypothesis that regions that harbor intact TEs (which are difficult to resolve in assemblies) are frequent targets of *trans*-regulation by the supergene. Regardless of genome build used, DAPs were retained on every chromosome and universally exhibited an enrichment of increased accessibility in *Sb* (Fig. 3d; Fig. S2; 75.87%, 89.01, and 89.75% of all DAPs mapped to chromosomes or scaffolds other than chromosome 16 in builds *Sb* v2, *SB* v2, and *SB* v3, respectively).

The variation in DAP distributions we observed across chromosomes and unplaced scaffolds between reference genome builds are consistent with an evolutionary history of *Sb* that includes some combination of translocation and copy number expansion. Nevertheless, many DEGs and DAPs that exhibit a particularly strong bias towards higher expression and chromatin accessibility in individuals carrying the *Sb* variant of the social chromosome do not exhibit evidence of supergene-linked CNV (Fig. 2b; (Fontana et al. 2020)). Overall, our analyses of gene expression, chromatin accessibility, CNVs, and intact TEs suggest the *Sb* supergene exhibits *trans*-regulatory effects that are influenced by a combination of copy number variation, gene regulatory evolution, and potentially, effects of TE density on *trans*-regulatory activation of host genes.

### Transcription factors may be drivers of global effects of the social chromosome

Based on the widespread effects of social chromosome genotype on chromatin accessibility and gene activity in the *S. invicta* genome (Fig. 1), we hypothesized the regulation of some transcription factors (TFs) may be influenced by social chromosome evolution. Indeed, examination of the orange gene co-expression module revealed that 9 of the 44 genes in the module are either orthologs of TFs or harbor DNA binding domains (Fig. 2b). Three distinct (non-CNV) uncharacterized proteins with Myb_SANT_like DNA binding domains are included in this total (described in relation to HARBI1 genes above), as well as a zinc finger BED domain-containing protein-encoding gene and several uncharacterized genes with zinc finger protein domains (Fig. 2b). Despite the constitutive effects of the supergene on expression in the orange module, none of the putative TFs in the module exhibit evidence of CNV between SB and Sb (Fig. 2b). These putative transcription factors nevertheless stand to play a particularly important role in *trans*-regulatory effects of the social chromosome.

We further identified eight putative TFs with evidence of *Sb*-linked CNV (Fontana et al. 2020) that lack constitutive variation in expression by supergene genotype (Table 2). Through context-specific regulation, these CNVs may convey direct *trans*-regulatory effects of the social chromosome, including in early developmental stages and specific tissues that have yet to be examined. Among the transcription factors exhibiting CNV between *Sb* and *SB*, four were also found to exhibit differential expression in one of two examined tissues of adult workers or queens between *SB/SB* and *SB/Sb* genotypes and four are near DAPs between *SB/Sb* and *SB/SB* worker brains (Table 2). Among orthologs of these genes, perturbation of both *cnc* (Deng and Kerppola 2013) and *btsz* (Serano and Rubin 2003) has been shown to influence body size at pupation in *Drosophila. S. invicta* orthologs of *cnc* and *btsz* thus represent excellent candidate factors for mediating effects of supergene genotype on worker size and queen determination (Buechel et al. 2014).

**Table 2.**
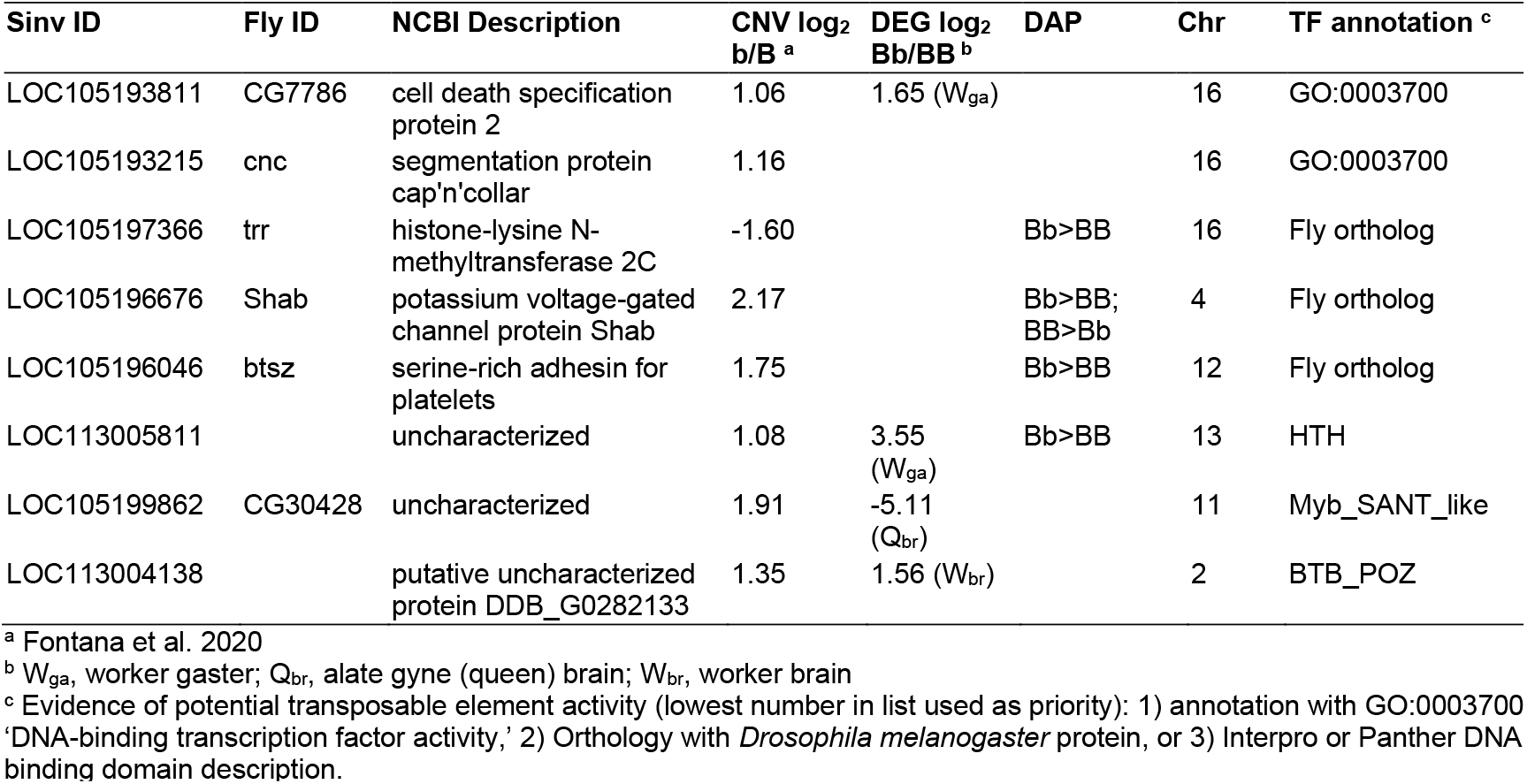
Transcription factors with social chromosome-linked copy number variation.

### Copy number variation and enhancer evolution act in concert to shape the expression of a supergene

To assess whether the social chromosome supergene and associated genotypic differentiation between *SB* and *Sb* alters transcriptional enhancer activity, we employed STARR-seq, a high-throughput reporter assay that quantifies the enhancer activity of randomly sheared genomic fragments (Arnold et al. 2013; Arnold et al. 2014; Jones et al. 2024). Using *Drosophila melanogaster* embryonic S2 cells, we identified 23,616 regions of the *S. invicta* genome with significant enhancer activity in either genotype (pools of either *SB* or *Sb* individuals, see Methods; (Jones et al. 2024)), 11,670 of which were active in both genotypes (Table S13). Enhancers were located primarily within introns (51.8%), promoter regions (13.9%) and transcription start sites (TSSs, 200bp flanking regions; 11.8%) of 5293 genes (Fig. S4). Supportive of the potential regulatory function of these regions, STARR-seq peaks were significantly more likely to overlap accessible chromatin peaks according to ATAC-seq than expected by chance (Fig. S5; permutations, 1.8-fold enrichment, p<0.005).

Unlike our finding of consistently more accessible chromatin in *Sb*-containing individuals, we did not see the same bias for enhancer activity when comparing SB and Sb DNA pools (i.e., the numbers of regions with biased activity toward *Sb* or *SB* pools were similar; Fig. S6, Table S13). This apparent discordance between regulatory activity of *Sb* inferred from ATAC-seq and STARR-seq suggests that while underlying sequence variation between genotypes influences enhancer activity, it is critical to also consider epigenetic context to understand how sequence variation leads to different phenotypes. For example, overlapping our enhancer set with DAPs suggests that approximately 12% of differentially accessible chromatin regions are enhancers (174/1454 DAPs overlap enhancer regions identified by STARR-seq). These peaks may be especially informative for understanding how supergene-linked variation influences gene regulation. Genes proximal to all enhancers with *SB*-biased activity (n=3378) were enriched for 81 GO terms, including terms related to signal transduction, cell adhesion, metabolism, axon guidance and synapse organization (Table S14, Fig. S7), while genes proximal to all *Sb*-biased enhancers (n=4079) were enriched for 61 GO terms related to transcriptional and protein biosynthetic processes, as well as terms related to neuronal processes (Table S15, Fig. S8).

Importantly, while sequence-based differences in enhancer activity are revealed by STARR-seq, the availability of upstream regulatory proteins and chromatin modifications will influence how this enhancer activity is realized in vivo. In addition, the pools of DNA used for STARR-seq included random genetic variation unrelated to social forms. Thus, we leveraged existing population genetics data (Wang et al. 2013; Martinez-Ruiz et al. 2020) (Table S16) to identify the location of variable loci between *SB* and *Sb* and tested for associations of these alleles with enhancer variation. First, we identified 283,725 loci with variation in the published datasets, which included US ((Wang et al. 2013); n=7 SB and Sb) and Argentine ((Martinez-Ruiz et al. 2020); n=13 SB and Sb) populations of *S. invicta*. Of the identified variants, nearly 20% (55116/283725) were located within enhancer peaks. While most of these variants were not fixed between *SB* and *Sb* genotypes, we identified 797 loci with differences in allele frequencies >80% between genotypes, 160 of which were divergent in both populations. Of those sites, 94 (of 797) were located within enhancer regions, providing a list of candidate loci that may influence regulatory activity. To assess the putative function of these loci, we tested for predictive effects of these SNPs on TF binding using the FIMO tool of MEME suite and the JASPAR 2024 set of core insect TF motifs (Rauluseviciute et al. 2024). Considering a region centered on the SNP with 50bp flanks, 49 of the 94 candidate regions had significant effects on at least one TF binding motif (Table S17). Motifs with the greatest numbers of predicted effects included those for Mothers against dpp (Mad), Clamp, Odd paired (Opa), and Dorsal (Dl), each of which had changes in predicted binding at 4 loci (Table S17). These TFs have numerous known roles through development (Jenkins et al. 2022) but Mad, Clamp, and Opa have caste-biased occupancy in honey bees (Lowe et al. 2022) and Clamp has also been associated with the major worker phenotype in ants (Glastad et al. 2020), suggesting these TFs may be frequently coopted for social-related phenotypes during evolution. It is notable in this context that *SB/Sb* larvae are more likely than *SB/SB* larvae to develop into large (major) workers and gynes (Buechel et al. 2014).

Of the predicted motif effects of divergence between *Sb* and *SB*, 17 were consistent with differences in enhancer activity between *SB* and *Sb* sequences based on STARR-seq, including 6 SNPs with fixed differences across both populations (US and Argentina) and significant correlations between allele frequencies at these loci and enhancer activity. These 6 variants are located within the supergene region in enhancers upstream or intronic to 4 genes, including *LOC105195710, cytoplasmic polyadenylation element-binding protein 2, cell death specification protein 2* and *nucleolysin TIAR* (Fig. 4). Future work should investigate the functional effects of these variants on TF binding as well as gene expression of putatively regulated genes.

**Figure 4.**
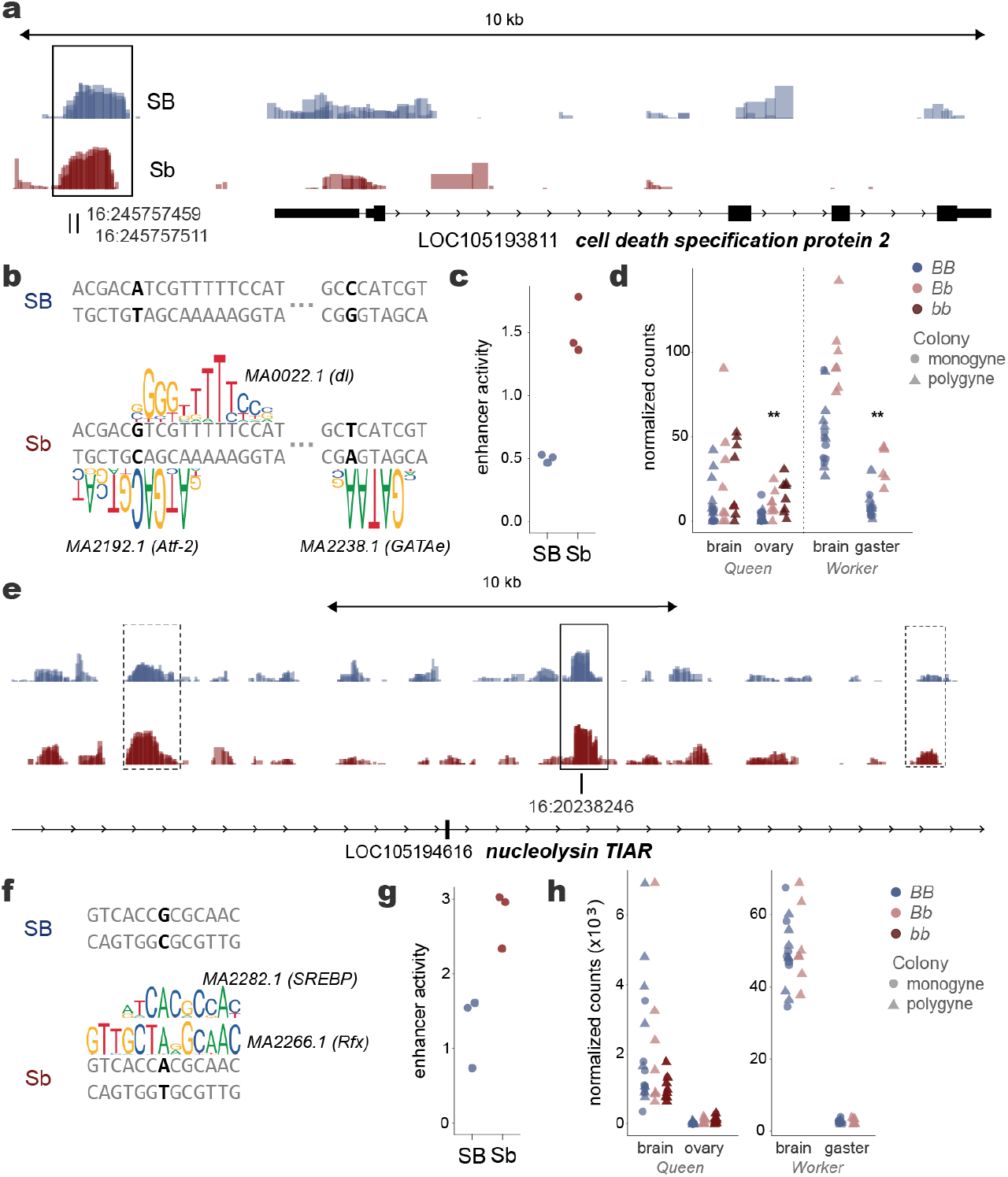
Differential enhancer activity coinciding with substitutions that affect predicted transcription factor binding. **(a)** Enhancer activity (displayed as bigwig track of log2 normalized ratios between mRNA and input DNA per replicate) according to STARR-seq, with increased enhancer activity in *Sb* relative to *SB* for an enhancer (within rectangle) in the upstream region of the transcription factor LOC105193811 *cell death specification protein 2* and **(b)** effects of two substitutions between *Sb* and *SB* on predicted TF binding within the enhancer. **(c)** Increased enhancer activity in *Sb* based on STARR-seq mirrors the increase in predicted binding of TFs. **(d)** Gene expression of *Sb*-carrying individuals is elevated in alate gyne (queen) ovaries and worker gasters, based on previously published data (Arsenault et al. 2020; Arsenault et al. 2023). **(e)** Enhancer activity according to STARR-seq, wherein 3 intronic enhancers of LOC105194616 *nucleolysin TIAR* show *Sb*-biased activity, including an enhancer (solid rectangle) with a substitution in *Sb* relative to *SB*. **(f)** Effects of a substitution between *Sb* and *SB* on predicted TF binding within the enhancer. **(g)** Increased activity of enhancer in *Sb* based on STARR-seq. **(h)** Gene expression is not significantly different by genotype in queens or workers for this gene.

Our integrative analysis of copy number variation, enhancer activity, and sequence substitutions between *SB* and *Sb* provides insight into how copy number variation may interact with enhancer evolution to fine tune gene expression. Among the putative TFs with copy number variation and orthologs of known function, we observed a significant differentially active enhancer (DAE), with higher activity in *Sb* than in *SB* in the upstream promoter region of LOC105193811 *cell death specification protein 2* (Fig. 4a; log2(*b/B*) 1.03, q-val=0.0003), a TF encoding gene that also has an extra copy in *Sb* (Fontana et al. 2020). Within this DAE we observed two substitutions between *SB* and *Sb*. Both variants affect the predicted binding affinity of TF binding motifs according to FIMO (Table S17), all of which lead to increased predicted binding of the *Sb* versus *SB* sequence (Fig. 4b), providing a putative mechanism for the increased enhancer activity (Fig. 4c) of the *Sb* sequence. These data suggest *cis*-regulatory evolution has acted to increase expression of *cell death specification protein 2* in *Sb*-carrying individuals. In this case, *cis*-regulatory evolution appears to have acted in concert with structural evolution to increase expression of a putative transcription factor. As further evidence of regulatory significance, we note that *cell death specification protein 2* is differentially expressed between *SB/SB* and *SB/Sb* in worker gasters (log2 ratio 1.65, FDR < 0.01) and in alternative homozygotes (*SB/SB* vs. *Sb/Sb*) in queen ovaries (log2 ratio 3.88, FDR < 0.01; (Arsenault et al. 2020)) (Fig. 4d).

Our analysis also revealed a potential example of evolutionary gene dosage compensation in the context of copy number variation for LOC105194616 *nucleolysin TIAR*, which encodes a protein that likely binds RNA and targets lysosomes to help induce apoptosis (Kawakami et al. 1992). This gene exhibits two copies in the ancestral *SB* (also *LOC105196096*) but has lost a copy in *Sb* (Table S7; (Fontana et al. 2020)) and is not differentially expressed between *SB/Sb* and *SB/SB* samples. We observed three intronic enhancers of *LOC105194616*, one of which contained a fixed variant and showed higher activity in *Sb* compared with *SB* (Fig. 4e; log2(b/B) 1.48, q-val=0.0003). The substitution between *SB* and *Sb* in this DAE affects the predicted binding affinity of two different TFs according to FIMO, each of which has increased binding of the *Sb* versus *SB* sequence (Fig. 4f, Table S17), providing a putative mechanism for the increased enhancer activity of the *Sb* sequence (Fig. 4g). In this case, *cis*-regulatory evolution appears to have acted in concert with structural evolution to maintain the correct dosage of *nucleolysin TIAR* (i.e., this gene is not differentially expressed between genotypes despite copy number variation; Fig. 4h).

## Conclusion

The fire ant supergene is evolutionarily young but structurally diverse, providing an ideal model to examine how regulatory sequence divergence, CNVs, and TEs interact to influence genome regulation. Our investigation revealed the derived supergene haplotype is associated with widespread increases in chromatin accessibility in genomic regions containing intact TEs. Thus, a relationship between chromatin accessibility and transcription factor binding sites associated with transposon activation could potentially contribute to the observed prevalence of *Sb*-activating effects on gene expression. Such a scenario would also be consistent with our finding of *Sb trans*-upregulation of genes domesticated from an ancestral transposon. Importantly, our results highlight that multiple orthologs of functionally annotated *Drosophila* TFs exhibit social chromosome linked CNVs in *S. invicta*. Enhancer activity differences by supergene haplotype revealed that regulatory element sequence evolution can amplify or mitigate effects of such CNVs. Overall, this research highlights that CNVs and regulatory sequences evolve in concert to shape the expression of supergene encoded factors with *trans*-regulatory links to the remainder of the genome.

## Methods

### Gene expression

#### Data sources

We compiled RNA-seq data from 122 libraries (Table S1), detailed as follows: Chandra et al. (2018) contributed 20 brain libraries split among workers (4 *SB/SB*; 6 *SB/Sb*), gynes (2 *SB/SB*; 5 *SB/Sb*), and reproductive queens (3 *SB/Sb*) from lab-reared colonies originally collected in Gainesville, FL. Arsenault et al. (2020) provided 63 brain and ovary libraries from 32 individual gynes (referred to as queens in main text) collected while embarking on spring mating flights around Athens, Georgia, USA, with 8 each of monogyne brain and ovary (all *SB/SB*) and 8 polygyne trios in both tissues (*SB/SB, SB/Sb, Sb/Sb*). Arsenault et al. (2023) added 39 libraries from individual monogyne workers (7 brain, 6 gaster; all *SB/SB*) and polygyne workers (7 brain and 6 gaster libraries each of *SB/SB* and *SB/Sb*) collected from age and size matched two-week old adult workers from lab reared colonies originally collected around Gainesville, Florida, USA. For further RNA-seq data generation details, see the source publications.

#### Quality control and read alignment

We trimmed reads and performed quality control using Trim Galore! v0.6.7. We then used STAR v2.7.10b (Dobin et al. 2013) to align reads using the 2-pass alignment procedure to the UNIL 3.0 genome assembly and associated annotation generated from an *SB* haploid male (GCF_016802725.1) (Helleu et al. 2022). After alignment, single-end worker libraries (Chandra et al. 2018) had 6-15 million, single-end gyne libraries (Arsenault et al. 2020) had 11-32 million uniquely mapped reads, and paired-end worker libraries (Arsenault et al. 2023) had 32-163 million. We loaded quantified, gene-level read counts found using featureCounts (Liao et al. 2014) into R Statistical Software (R_Core_Team) for subsequent analyses.

#### Differential expression analysis

We used DESeq2 (Love et al. 2014) to perform separate differential expression analyses for gyne data from Arsenault et al. (2020) and worker data from Arsenault et al. (2023). First, we performed principal component and hierarchical clustering for each data set comparison separately using the normalized counts. To visualize expression variance and relationships between libraries, HCA was performed with the R package pheatmap (Kolde and Kolde 2018) using the “ward D2” clustering method. All differentially expressed genes were called using a 5% false-discovery rate threshold (FDR<0.05). All pairwise comparisons of gene expression are provided in Table S2.

#### Weighted gene co-expression network analysis

Normalized counts from all 122 libraries were converted to an expression matrix for WGCNA (v.1.72-5) (Langfelder and Horvath 2008). We applied a low expression filter to remove genes that had a normalized count value less than 10 for more than 90% (109/122) of the libraries. We then performed a variance stabilizing transformation using varianceStabilizingTransformation from DESeq2 on the filtered raw expression data. To visualize expression variance and relationships between all libraries, HCA was performed with the R package pheatmap (Kolde and Kolde 2018) using the “ward D2” clustering method. We also performed a PCA using all samples (Fig. S9).

Adjacency matrices were raised to a soft power threshold of 24. This was empirically determined based on a measure of R^2 scale-free topology model fit that maximized and plateaued over 0.9. The soft-power thresholded adjacency matrices were converted into a topological overlap matrix (TOM) and a topological dissimilarity matrix (1-TOM). We then performed agglomerative hierarchical clustering using the average linking method on the TOM dissimilarity matrix. Gene modules were defined from the resulting clustering tree (Fig. S10), and branches were cut using the hybrid dynamic tree cutting function: the module detection sensitivity (deepSplit) was set to 2, minimum module size 20, and the cut height for module merging set to 0.15 (modules whose eigengenes were correlated above 0.85 were merged). This yielded 26 consensus modules which were each assigned a color label (Table S8). For each gene module, a summary measure (module eigengene) was computed as the first principal component of the module expression profiles.

### Chromatin accessibility

#### Worker brain cell suspension for ATAC-seq

Individual adult workers of unknown age were sampled from a lab-reared colony initially collected around Athens, Georgia, USA. Brains of *S. invicta* workers were dissected under an Olympus SZ61 microscope and stored individually in 1.5mL microcentrifuge tubes containing 50μL Hank’s Balanced Salt Solution (HBSS) (H6648-500ML, Sigma). Tubes were kept on ice to preserve the cells. 950μL of a solution of Papain (LK003178, Worthington Biochemical) mixed with 5mL HBSS was added to each tube, and the tubes were incubated in a thermal mixer (13687720, Thermo Scientific) for 15 min at 25°C, with 900 RPM rotation/mixing. After incubation, the Papain-HBSS buffer was removed and brains were resuspended in 200μL phosphate buffered saline (PBS) (806544-500ML, Sigma) + protease inhibitor complex (PIC) (Roche 4693132001). Each brain was then aspirated in 100μL PBS-PIC, transferred to a glass dounce, and gently homogenized with two pestles. The homogenate was transferred to a new 1.5μL microcentrifuge tube; the dounce and pestles were rinsed in the remaining 100μL PBS-PIC, and the rinsate was added to the new tube, to increase recovery.

#### ATAC-seq transposition reaction

Cells were pelleted at 500 RCF (0.5×1000) at 4°C for 5 min, after which PBS-PIC supernatant was removed, and 50μL of cold ATAC-Resuspension Buffer (RSB) (500μL 1M Tris-HCL [15567-027, Invitrogen], 100μL 5M NaCl [AM9759, Ambion/Thermo], 150μL 1M MgCl2[AM9530G, Ambion/Thermo], 49.25mL sterile H20) containing 0.1% NP40 substitute (2127-50, BioVision), 0.1% Tween-20 (11332465001, Sigma/Roche), and 0.01% Digitonin (G9441, Promega) was added to lyse cells. The RSB containing detergents was gently pipetted up and down three times, and the mixture was incubated on ice for 3 min. To inhibit lysis, 1mL of cold ATAC-RSB containing only 0.1% Tween-20 was then added, and tubes were inverted 3 times. Nuclei were pelleted at 500 RCF for 10 min at 4°C.

All supernatant was aspirated, taking care to avoid the cell pellets, and the pellets were resuspended in 50μL of transposition mixture (25μL 2X TD buffer, 2.5μL transposase (100nM final), 16.5μL PBS, 0.5μL 1% Digitonin, 0.5μL 10% Tween-20, 5μL H20) by pipetting up and down 6 times. The reaction was incubated at 37°C for 30 min in the thermal mixer described above with 1000 RPM mixing. Reactions were cleaned using a DNA Clean and Concentrator-5 Kit (D4014, Zymo), using 250μL DNA Binding Buffer. DNA was eluted in 25μL elution buffer and stored at −20°C until amplification.

#### Amplification of transposed fragments and sequencing

Transposed fragments were amplified using the following PCR recipe: 25μL 2X NEBNext® High-Fidelity 2X Master Mix (M0541L, NEB), 2.5μL of 25μM i5 primer, 2.5μL of 25μM i7 primer, and 20μL transposed/cleaned-up reaction. Different i7 and i5 primer combinations were used for each reaction. The thermocycler conditions are as follows: 72°C for 5 min, 98°C for 30 s, followed by 9 cycles of 98°C for 10 s, 63°C for 30 s, and 72°C for 1 min; final hold at 4°C. PCR products were purified with a DNA Clean and Concentrator-5 Kit (D4014, Zymo) using 250μL DNA Binding Buffer and eluted in 25μL sterile H20.

Fragment analysis on a bioanalyzer supported nucleosomal banding patterns for libraries selected for sequencing (Fig. S11). ATAC-seq libraries were pooled and sequenced on an Illumina NextSeq 2000 with 50 bp pairedend reads (Table S3). After filtering, an average of 121 million read pairs remained for each library (range: 108-142 million), 97% of which were mapped in proper pairs to the assembly with duplication rates of 24-27%. Additional information on sequencing coverage and alignment is in Table S3. Mean genomic coverage across samples ranged from 20.0-26.9 reads per bp (Fig. S12).

#### ATAC-seq analyses

FastQC (Andrews 2010) was used to examine data quality and raw reads were trimmed to remove low-quality bases and sequencing adapters with fastp v0.20.1 (--cut_right, --cut_right_window_size 4, --cut_right_mean_quality 15) (Chen et al. 2018). Trimmed reads were mapped to the *S. invicta* genome assembly UNIL_Sinv_3.0 with BWA v0.7.17 (bwa mem –M) (Li 2013) and sorted by read name using samtools v1.3 (samtools sort –n) (Danecek et al. 2021). Genrich v0.6.1 (https://github.com/jsh58/Genrich) was used to call peaks on each sample group (*BB* and *Bb*, including all samples of each type in the respective peak calling runs) in ATAC-seq mode with removal of PCR duplicates (Genrich –j –y –r). Merging Genrich peaks from both sample groups resulted in 33,619 total peaks, with an average of 31.4% of reads mapping to this peakset (Fraction of Reads in Peaks, or FRiP, ranged from .3015 to .3307; Table S3). These peaks were assigned to gene features using the publicly available RefSeq annotation GCF_016802725.1_UNIL_Sinv_ 3.0_genomic.gff, with TSS flanks defined as the TSS +/−200 bp, promoters defined as 0-5kb upstream of gene start, upstream regions defined as 0-10kb upstream of gene start, and downstream regions defined as 0-10kb downstream of gene start. Exons and introns were extracted from the RefSeq annotation. Peaks that did not overlap any of these features were annotated as intergenic, and peaks that overlapped multiple regions were assigned in the following priority: TSS flank, promoter, intron, upstream, downstream, exon, intergenic. The function overlapPermTest in the R package regioneR (Gel et al. 2016) was used to assess whether the overlap of peaks with particular features was more than expected by chance. ATAC-seq peaks were enriched for the following features: TSS flanks, promoters, upstream regions, downstream regions and exons (Fig. S13). In contrast, ATAC-seq peaks overlapped introns and intergenic regions less often than expected by chance (Fig. S13).

Differentially accessible peaks (DAPs, FDR<0.1; Table S4; Fig. S14) were identified using DiffBind v3.12.0 (https://bioconductor.org/packages/release/bioc/html/DiffBind.html) with default DESeq2 parameters using deduplicated BAM files (produced with picardtools v2.22.1 [http://broadinstitute.github.io/picard] following coordinate-sorting of BAMs by samtools) and Genrich called peaks as input (using group-specific Genrich peak calls for each respective sample type). Peaks were annotated as detailed above with both with the nearest priority-assigned gene and to all genes within 10kb of either end of the peak.

### Enhancer activity

#### Generation of STARR-seq plasmid libraries

Genomic DNA was isolated from individual male progeny of a lab-reared isolated *S. invicta* polygyne queen and a polygyne colony containing multiple queens, both collected around Athens, Georgia, USA, using a Zymo Quick-DNA Tissue/Insect Microprep Kit (Zymo Research cat. no. D6015). Extracted DNA from each individual male was genotyped twice at *Gp-9* to determine whether they were hemizygous or homozygous at the *Gp-9* locus, resulting in 9 males with *SB* and 10 males with *Sb*. These *SB* and *Sb* samples were pooled to obtain sufficient material to generate *SB*- and *Sb*-specific STARRseq plasmid libraries. Additional details on individuals used for each genotype are in Table S18. A pool of 5 ug from each genotype was sonicated in a microTUBE with AFA fiber (cat. no. 520045) using a Covaris LE220 and the following parameters: 45 sec sonication time, 450W peak incident power, 15% duty factor, 200 cycles per burst. Sheared DNA was run on a 1% agarose gel and 400-750bp fragments were size-selected via excision under blue light. Size-selected DNA was purified using a Gel DNA Recovery Kit (Zymo Research cat. no. D4008), followed by an additional purification with the QIAquick PCR Purification Kit (Qiagen cat. no. 28104). Size-selected DNA was quantified using a Qubit dsDNA HS Assay Kit (Invitrogen cat. no. Q32854) on a Qubit4 Fluorometer.

Illumina-compatible adapters were ligated to size-selected genomic DNA using the NEBNext Ultra II End Repair Module (NEB E7546L) with 1 ug fragmented DNA per genotype. Adapter-ligated DNA libraries were then cleaned with a 1.8x volume ratio of AMPure XP Reagent beads (Beckman Coulter cat. no. A63881) to sample following manufacturer protocols for PCR Purification, then cleaned a second time with 0.8x bead to sample volume ratio. Adapter-ligated DNA libraries were then amplified in 5 separate reactions for 5 cycles (PCR conditions: initial denaturation of 98C for 45 sec, then 5 cycles of 1) denaturation: 98C for 15 sec, 2) annealing: 65C for 30 sec, 3) elongation: 72C for 45 sec, and a final elongation at 72C for 60 sec) each using in_fusion_F (TAGAGCATGCACCGGACACTCTTTCCCTACACGA CGCTCTTCCGATCT) and in_fusion_R (GGCCGAATTCGTCGAGTGACTGGAGTTCAGACGT GTGCTCTTCCGATCT) primers at 10 uM and 2x KAPA HiFi HotStart Ready Mix (Roche cat. no. KK2601). PCR reactions were pooled for each genotype and cleaned with 0.8x bead to sample volume ratio with AMPure XP Reagent beads (Beckman Coulter cat. no. A63881) followed by an additional purification using the QIAquick PCR Purification Kit (Qiagen cat. no. 28104). Libraries were quantified using a Qubit dsDNA HS Assay Kit (Invitrogen cat. no. Q32854) on a Qubit4 Fluorometer and average sizes of each adapter-ligated, amplified library was determined with Agilent High Sensitivity DNA reagents on an Agilent 4200 TapeStation (Agilent cat. Nos. 5067-5592, 5067-5593, 5067-5594).

pSTARR-seq_fly was a gift from Alexander Stark (Addgene plasmid #71499; http://n2t.net/addgene:71499; RRID:Addgene_71499) (Arnold et al., 2013). The pSTARR-seq_fly reporter vector was digested with AgeI-HF (NEB cat. no. R3552S) and SalI-HF (NEB cat. no. R3138S) restriction enzymes with 250 units of each and 25 ug vector per reaction, incubated at 37C for 2h followed by a 20 min heat inactivation at 65C. Digested products were run on a 1% agarose gel and linearized vector was selected via excision under blue light and purified using a Gel DNA Recovery Kit (Zymo Research cat. no. D4008). Eluates from gel extraction were purified using a 1.8x bead cleanup with AMPure XP beads (Beckman Coulter cat. no. A63881). Adapter-ligated DNA libraries were cloned into purified pSTARR-seq_fly using a 2:1 molar ratio of insert (size determined via TapeStation) to plasmid (~4125bp). Two cloning reactions were conducted for each genotype using 1 ug digested plasmid, the appropriate amount of PCR amplified, adapter-ligated DNA library to have a 2x molar excess insert to plasmid, and 10 ul 5x In-Fusion HD Enzyme Premix (Takara cat. no. 638910) in a total volume of 50 ul. Reactions were incubated for 15 min at 50C, then 200 ul EB was added. Next, 25 ul 3M NaAc (pH 5.2) and 2 ul Pellet Paint Co-precipitant (Millipore cat. no. 69049), were added to each reaction, vortexing between each addition. Finally, 750 ul ice-cold 100% EtOH was added, samples were vortexed, then incubated at −20C for 16h. Samples were centrifuged for 15 min at full speed and 4C, vortexed, centrifuged again for 15 min, then supernatant was carefully aspirated. Cloned DNA pellets were washed 3 times with 750 ul ice-cold 70% EtOH, mixing each time by inversion. Cloned DNA pellets were again centrifuged for 15 min at full speed and 4C, supernatant aspirated, then pellets dried for 30 sec at 37C then further at room temperature until dry. Each pellet was resuspended in 12.5 ul EB and incubated for 3h at −80C prior to transfer to −20C for storage until transformation.

Cloned DNA reactions were transformed into electrocompetent MegaX DH10B cells (ThermoFisher cat. no. C640003) using 150 ul cells for each clone (two clones per genotype) split across two Gene Pulser Electroporation Cuvettes (0.1 cm gap, Bio-Rad cat. no. 1652089) and the Bio-Rad Gene Pulser Xcell system with the following electroporation conditions: 2 kV, 25 uF, 200 ohms. Immediately after electroporation, 500 ul of pre-warmed recovery medium was added and cells were transferred to round bottom tubes with an additional 4 ml warm recovery media. Transformed bacteria were incubated for 1h at 37C and 225 rpm, then each transformation reaction was added to 300 ml warm LB+ampicillin (100 ug/ml) in 2L flasks and incubated for 12h while shaking at 200 rpm at 37C. Cells were harvested via centrifugation and plasmids were purified using the ZymoPure II Plasmid Maxiprep Kit (Zymo Research cat. no. D4203) with a maximum of 75 ml culture per column and eluted in water heated to 50C prior to elution. Plasmids were pooled within each clone and then across clones from the same genotype, resulting in one clone library per genotype (i.e., *SB* library and *Sb* library).

#### Transfection of Drosophila cells with STARR-seq plasmid libraries

Three replicate flasks were seeded and transfected for each genotype (*SB* or *Sb*). *Drosophila* S2 cells (S2-DRSC; DGRC Stock 181; https://dgrc.bio.indiana.edu//stock/181; RRID:CVCL_Z992, (Schneider, 1972)) were cultured in M3 media (Sigma-Aldrich cat. no. S8398) supplemented with BactoPeptone (BD Biosciences #211677), yeast extract (Sigma-Aldrich Y1000), 10% heat-inactivated fetal bovine serum (Gibco cat. no. 10437-010) and 1% Penicillin/Streptomycin solution (ThermoFisher cat. no. 15140122) at 25C using standard cell culturing protocols. 24 hours prior to transfection, cells were split, washed, counted, and seeded at a density of 27 million cells in 15 ml media per T75 flask, with 3 flasks seeded per genotype. Effectene Transfection Reagent (Qiagen cat. no. 301427) was used to transfect 3 replicate flasks of cells per genotype. For each replicate, 12 ug plasmid clone library was diluted with Buffer EC to a total volume of 450 ul per flask, 96 ul enhancer was added and the reaction was vortexed briefly then incubated for 5 minutes. 150 ul effectene was added, the solution was mixed by pipetting up and down 5 times, then incubated for 10 minutes. One ml media was added to the complex, mixed by pipetting up and down twice, then added drop-wise onto the flask of cells. Flasks were swirled gently to ensure uniform distribution then returned to 25C incubator for 48 hours until harvest.

For harvesting, cells were gently pipetted to bring into suspension then centrifuged for 5 min at 350g. Cells were washed once with 10 ml 1X PBS. then incubated at 37C for 5 min in 2 ml M3+BPYE media containing 1 ml Turbo DNase (ThermoFisher cat. no. AM2239) per 36 ml. Cells were again pelleted via centrifugation, supernatant removed, then resuspended in 10 ml 1X PBS. An aliquot of 10% unlysed cells per flask was set aside for later plasmid extraction (pelleted and stored at −20C) and the remaining cells were pelleted and lysed in 2 ml RLT (Qiagen RNeasy Midi Kit cat. no. 75144) plus 20 ul 2-mercaptoethanol (Sigma cat. no. 60-24-2) then frozen at −80C.

#### Generation of input and STARR-seq libraries

For each of the six transfected flasks, both an input library (derived from fragment inserts of plasmid DNA purified from transfected cells) and a STARR library (derived from plasmid-derived mRNA) were generated. Plasmid DNA was purified from 10% of harvested cells per flask using a QIAprep Spin Miniprep Kit (Qiagen cat. no. 27104). Total RNA was extracted from 90% of harvested cells per flask using a Qiagen RNeasy Midi Kit (Qiagen cat. no. 75144). mRNA was isolated from 75 ug total RNA using Dynabeads Oligo(dT)25 (Invitrogen cat. no. 61005), followed by a DNase digestion with Turbo DNaseI (ThermoFisher cat. no. AM2239). RNA was cleaned with RNAClean XP beads (Beckman Coulter cat. no. A63987) using a 1.8x bead to sample volume ratio and reverse transcribed using SuperScriptIII (Invitrogen cat. no. 18080093) using a gene-specific primer (CTCATCAATGTATCTTATCATGTCTG). cDNA was purified using a 1.8x bead to sample volume of AMPure XP beads (Beckman Coulter cat. no. A63882) and used in a junction PCR to amplify only plasmid-derived mRNA with primers that span a synthetic intron of the pSTARR-seq_fly reporter vector. cDNA was used in this junction PCR with 15 cycles (PCR conditions: initial denaturation of 98C for 45 sec, then 15 cycles of 1) denaturation: 98C for 15 sec, 2) annealing: 65C for 30 sec, 3) elongation: 72C for 70 sec, and a final elongation at 72C for 60 sec) each using junction_F (TCGTGAGGCACTGGGCAG*G*T*G*T*C) and junction_R (CTTATCATGTCTGCTCGA*A*G*C) primers and 2x KAPA HiFi HotStart Ready Mix (Roche cat. no. KK2601). Junction PCR products were purified with a 0.8x bead to sample volume ratio with AMPure XP Reagent beads (Beckman Coulter cat. no. A63881). Either 10 ng plasmid DNA (input) or entire cleaned junction PCR products (STARR libraries) were used as input for a PCR to add indices for sequencing, and final libraries were quantified using the dsDNA HS Assay Kit (Invitrogen cat. no. Q32854) on a Qubit4 Fluorometer and average sizes of each library was determined with Agilent High Sensitivity DNA reagents on an Agilent 4200 TapeStation (Agilent cat. Nos. 5067-5592, 5067-5593, 5067-5594). Libraries were sequenced on one flowcell of a NovaSeq SP with 2×50nt paired-end reads at the Genomics Core Facility at Princeton University. Additional library information and accession numbers are in Table S18.

#### STARR-seq data processing and identification of enhancers

Raw FASTQ files were processed to remove low quality bases and adapter contamination using fastp (Chen et al. 2018) with default parameters. Processed FASTQ files were then aligned to the *S. invicta* genome (GCF_016802725.1) with bwa mem (Li 2013) and sorted with SAMtools (Danecek et al. 2021). Enhancer peaks were called with MACS2 (Zhang et al. 2008) on each replicate flask, with input libraries as controls and the following parameters: -f BAMPE, -g 3.8e8 –keep-dup all -q 0.05.

Peaks called on each flask (3 per genotype, 6 total) were concatenated, sorted and merged with BEDtools (Quinlan and Hall 2010), resulting in a set of 67,105 peaks. This consensus set was used with featureCounts from the Subread package (Liao et al. 2014) to count reads mapping to each peak region for all input and STARR libraries. The R packages limma (Ritchie et al. 2015) and edgeR (Robinson et al. 2009) were used to identify genomic regions with significant enhancer activity using the following model: ~genotype+librarytype:genotype. 23,616 peaks had significant positive enhancer activity in one or both genotypes and were retained. Enhancer peaks were assigned to gene features using the publicly available RefSeq annotation GCF_016802725.1_UNIL_Sinv_3.0_genomic.gff, with TSS flanks defined as the TSS +/−200 bp, promoters defined as 0-5kb upstream of gene start, upstream regions defined as 0-10kb upstream of gene start, and downstream regions defined as 0-10kb downstream of gene start. Exons and introns were extracted from the RefSeq annotation. Peaks that did not overlap any of these features were annotated as intergenic, and peaks that overlapped multiple regions were assigned in the following priority: TSS flank, promoter, intron, upstream, downstream, exon, intergenic. The function overlapPermTest in the R package regioneR (Gel et al. 2016) was used to assess whether the overlap of peaks with particular features was more than expected by chance.

Differentially active enhancers (DAEs) between SB and Sb were identified using the R packages limma (Ritchie et al. 2015) and edgeR (Robinson et al. 2009). Peaks were tested for significant interactions between genotype (SB and Sb) and library type (RNA or DNA) using the following model design: ~genotype+librarytype+genotype*librarytype. The qvalue package (Storey et al. 2024) was used to correct p-values for the interaction term, and peaks with q<0.05 were considered DAEs. For plotting purposes, enhancer strength was quantified as the log2 fold-change of normalized counts from mRNA (STARR libraries) relative to DNA input (input libraries) for each replicate.

### Annotation

#### Transcription factor motif and GO enrichment

To determine whether specific regulatory motifs are involved in supergene-mediated regulatory variation, we used HOMER (v4.10, (Heinz et al. 2010)) and the JASPAR2024 database of insect motifs (Rauluseviciute et al. 2024) to identify enriched TF motifs with ATAC-seq and STARR-seq peaks related to supergene genotype. Specifically, the findMotifsGenome.pl script was used with input files including differential ATAC-seq peaks (DAPs) and differential STARR-seq peaks (DAEs) related to supergene genotype. Results of HOMER and GO for DAPs and DAEs are provided in Table S5, S6, S14, and S15. GOATools (Klopfenstein et al. 2018) was used for Gene Ontology analyses of ATAC-seq and STARR-seq peak datasets. Visualization of GO enriched terms by semantic similarity (as in Fig. S5-S6) was performed using GO-Figure! (Reijnders and Waterhouse 2021).

#### Transcription factor orthologs

To identify a set of putative transcription factors in the *S. invicta* genome, we performed a one-way blastp querying the same fire ant genome used for mapping RNA-seq data (GCF_016802725.1) against the latest available *Drosophila melanogaster* genome (GCF_000001215.4). We identified the best hit based on multiple criteria, percent sequence identity, alignment length, E-value, and bitscore. This approach reduced our set of 14,064 hits to 9,368. We then used FlyBase to identify *D. melanogaster* genes annotated to ‘DNA-binding transcription factor activity’(GO:0003700). We used these FlyBase IDs to subset the *D. melanogaster* proteins and identify the *S. invicta* genes that encoded the *S. invicta* proteins found as best hits to *D. melanogaster* transcription factor protein. Of the 555 *D. melanogaster* genes annotated to ‘*DNA-binding transcription factor activity*’(GO:0003700), we identified 452 *S. invicta* orthologs which served as our set of putative DNA-binding transcription factors. For a more inclusive list of TFs, amino acid sequences of all protein-coding genes were first scanned with InterProScan (Jones et al. 2014) and all genes with a significant domain hit to a known DNA binding domain (Zf_C2H2, homeobox_like, HTH, WH, LIM_type, HMG_box, Zf_GATA, Pax, bHLH, BTB_POZ, POU, bZIP, p53_like,Zf_TFIIB_type, MADS_box, Znf_hrmn_rcpt) were treated as putative DNA-binding factors.

#### Copy number variants

We annotated genes in GCF_016802725.1_UNIL_Sinv_3.0 by their CNV status using a previously published set of CNV transcripts (Fontana et al. 2020). To map transcripts to the current assembly, we used GMAP (Wu and Watanabe 2005) to align all nr-sinv-ref.fasta transcripts, then overlapped predicted exons from GMAP to gene models. Hits with <90% sequence identify were removed, resulting in 25,094 transcripts with unique hits and 1,475 transcripts with multiple hits to 14,192 annotated genes in UNIL 3.0 (*SB*). Next, CNV log2 values (*Sb* vs. *SB*) from supplemental information of Fontana et al. (2020) were used to infer CNV direction and strength for each gene with a significant CNV transcript hit. Because many of the CNV transcripts map to more than one locus, the 201 CNV transcripts with at least one >90% identity to exons (of 260 reported CNVs in Fontana et al. 2020) resulted in CNV values for 267 total genes. In addition, some genes had multiple significant transcripts aligning to them; in this case, we selected the highest magnitude log2 fold-change value for that gene when visualizing CNV strength. CNV values for all genes with hits from significant transcripts are provided in Table S7.

#### Transposable elements

In this study, transposable element (TE) annotation was conducted using the Extensive de novo TE Annotator (EDTA) version 2.1.0 (Ou et al. 2019) For the reference genome assembly (GCF_016802725.1_UNIL_Sinv_3.0), the analysis utilized the genome file (GCF_016802725.1_UNIL_ Sinv_3.0_genomic.fna) and its corresponding coding sequences (CDS; GCF_016802725.1_UNIL_Sinv_3.0_ cds_from_genomic.fna). The EDTA analysis was executed with parameters to prevent file overwriting (--overwrite 0), enhance TE detection sensitivity (--sensitive 1), perform TE annotation (--anno 1), evaluate annotation quality (--evaluate 1), and force execution (--force 1). To evaluate TE content in earlier genome builds (GCA_009650705.1_Solenopsis_invicta_SB1.0 and GCA_010367695.1_ASM1036769v1), transcriptomic sequences (rna.fna) from the reference assembly were mapped to these genomes using GMAP-GSNAP v11.3.0 ((Wu et al. 2016)) with a minimum identity threshold of 90% (--min-identity=0.90) and options for generating gene-structured GFF3 output (--gff3-fasta-annotation=1 and -f gff3_gene). The resulting GFF3 files were then refined using the Another Gtf/Gff Analysis Toolkit (AGAT) v1.1.0 ((Dainat)). In particular, the script agat_sp_fix_features_locations_duplicated.pl was executed with the “--model 1,2,3” option to address various annotation scenarios. For instance, when multiple isoforms exhibited identical exon arrangements, redundant entries were culled by retaining only the transcript with the longest CDS. Similarly, if mRNA entries from distinct gene identifiers had the same exon structure but lacked CDS information, one duplicate was removed. In cases where transcripts from different genes had both matching exon and CDS patterns, only the transcript with the longest CDS was preserved. Moreover, when transcripts shared an identical exon structure yet differed in CDS configuration, the tool adjusted the untranslated regions to redefine mRNA and gene boundaries. Lastly, for overlapping mRNA entries with differing exon architectures, AGAT modified gene coordinates by trimming the UTRs. The corrected GFF3 files were subsequently processed using the agat_sp_extract_sequences.pl script to extract refined sequences, which then served as input for EDTA, ensuring a consistent TE annotation process across all genome builds.

#### Variant calling, filtering, and visualization

In this study, we utilized SAMtools v1.16.1 (Danecek et al. 2021) and Picard v2.27.5 (Broad_Institute 2019) in conjunction with GATK v4.4.0.0 (McKenna et al. 2010) for data preprocessing and variant calling. The GATK functions, including HaplotypeCaller with the -ERC GVCF mode, GenomicsDBImport, GenotypeGVCFs, and VariantFiltration, were used to identify and filter variants from both ATAC-seq and STARR-seq datasets. Filtering criteria applied during variant calling included thresholds for depth (DP < 10.0), quality (QUAL < 30.0), quality by depth (QD < 2.0), mapping quality (MQ < 40.0), FisherStrand (FS > 60.0), Strand Odds Ratio (SOR < 3.0), Mapping Quality Rank Sum Test (MQRankSum < −12.5), and Read Position Rank Sum Test (ReadPosRankSum < −8.0). Additionally, further filtering to select for biallelic sites was conducted using BCFtools v1.15.1 (Danecek et al. 2021) with parameters: -m2 -M2 -v snps. Subsequently, the filtered reads were converted into a variants table using the GATK VariantsToTable function with following parameters: -F CHROM -F POS -F TYPE -F REF -F ALT -GF AD. Variants were visualized using the VIVA VariantVisualization program (Tollefson et al. 2019), with the parameters: -x --avg_dp sample,variant -- save_remotely.

Previously published raw reads from fire ant samples collected in Georgia (PRJNA182127; (Martinez-Ruiz et al. 2020)) and Argentina (PRJNA542606; (Martinez-Ruiz et al. 2020)) were downloaded using the SRA-Toolkit v3.0.1 (fasterq-dump; https://github.com/ncbi/sra-tools/wiki/HowTo:-fasterq-dump). See Table S16 for sample details. Adapter removal was performed on the raw FASTQ files using fastp v0.23.2 ((Chen et al. 2018)), and the trimmed reads were mapped to the reference genome (GCF_016802725.1_UNIL_Sinv_3.0) using BWA v0.7.17 (mem; (Li 2013)). The resulting alignments were sorted with SAMtools, and variants were subsequently called using GATK with the same parameters described above.

#### Evaluation of SNP effects on predicted TF binding and enhancer activity

Fixed or nearly fixed variants between *SB* and *Sb* were determined using the above population genetics datasets including individuals from Argentina and Georgia. Variants of interest were defined as follows for each population: fixed variants had reference allele frequencies of 1 for either *SB* or *Sb* and 0 for the other genotype, while nearly fixed variants had reference allele frequencies of at least 0.8 for either *SB* or *Sb* and less than 0.2 for the other genotype. Of 283,725 total SNPs called across Argentinian and Georgian populations, 679 were nearly fixed between *SB* and *Sb* in Georgia samples, while 331 SNPs were nearly fixed between *SB* and *Sb* in Argentina samples, with 160 SNPs common to both (Table S19). For completely fixed variants, we identified 573 in Georgia and 183 in Argentina between *SB* and *Sb* individuals, 60 of which were common to both (Table S19).

For each SNP, two 101-bp regions (one for each allele) centered on the SNP of interest were scanned with FIMO (Grant et al. 2011) using the JASPAR2024 database of nonredundant insect motifs (Rauluseviciute et al. 2024). A SNP was considered to have an effect on predicted binding for a given motif if only one allele had a significant (p < 0.0001) match for that motif and/or if the ratio of FIMO scores between the two alleles was >1.5, as in (Kapheim et al. 2020; Jones et al. 2024). Results of FIMO are provided in Table S17.

## Data Availability

Sequencing data is deposited in NCBI’s Sequencing Read Archive under accession PRJNA1071646.

## Author Contributions

BMJ, AHW, SDK and BGH designed and conceptualized the study. BMJ and SK generated new data, and data curation was handled by BMJ, AHW, and MAC. Data analysis was performed by BMJ, AHW, MAC, KMG, and BGH. BMJ and BGH designed figures and wrote the initial draft of the manuscript. BGH supervised the work. All authors reviewed and edited the final manuscript.

## Acknowledgements

This work was supported by US NSF grants 1754476 to SK and BH, 1755130 to BH, 2105033 to MG, and support to BMJ from the University of Kentucky. We thank the Drosophila Genomics Resource Center (NIH Grant 2P40OD010949) for maintenance of and access to cell lines, the University of Georgia Genomics and Bioinformatics Core for sequencing of ATAC-seq libraries, and the Genomics Core Facility at the Lewis-Sigler Institute for Integrative Genomics at Princeton University for sequencing of STARR-seq libraries.

## Conflict of Interest

The authors declare no conflicts of interest.

## Notes

### Competing Interest Statement

The authors have declared no competing interest.

