## Supplemental Figures for "The fire ant social chromosome exerts a major influence on genome regulation"

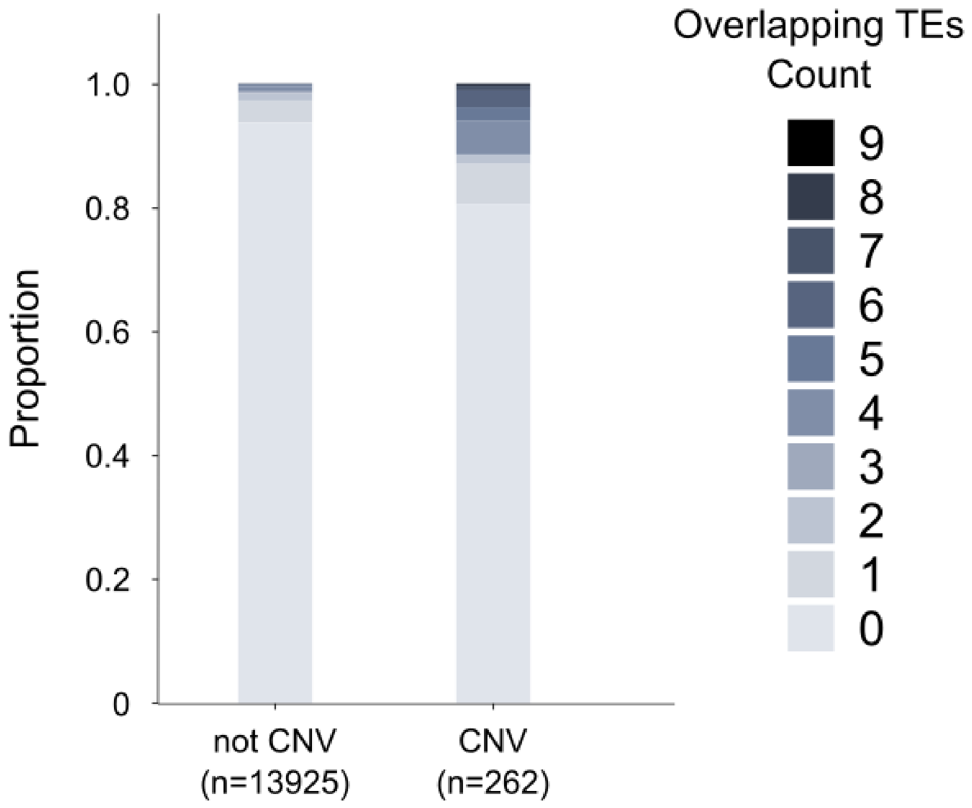


**Figure S1. CNVs are more likely to overlap intact TEs than non-CNVs**

Across the genome, genes with evidence of *Sb*-linked copy number variation (CNV) (Fontana et al. 2020) exhibit increased overlap with intact TEs as compared to non-CNVs. Only 6.41% of non-CNVs overlap with at least one TE, while 19.47% of CNVs overlap at least one TE. Fisher's Exact test, odds ration=3.6, p=1.3e-12. CNV information provided in Table S7.

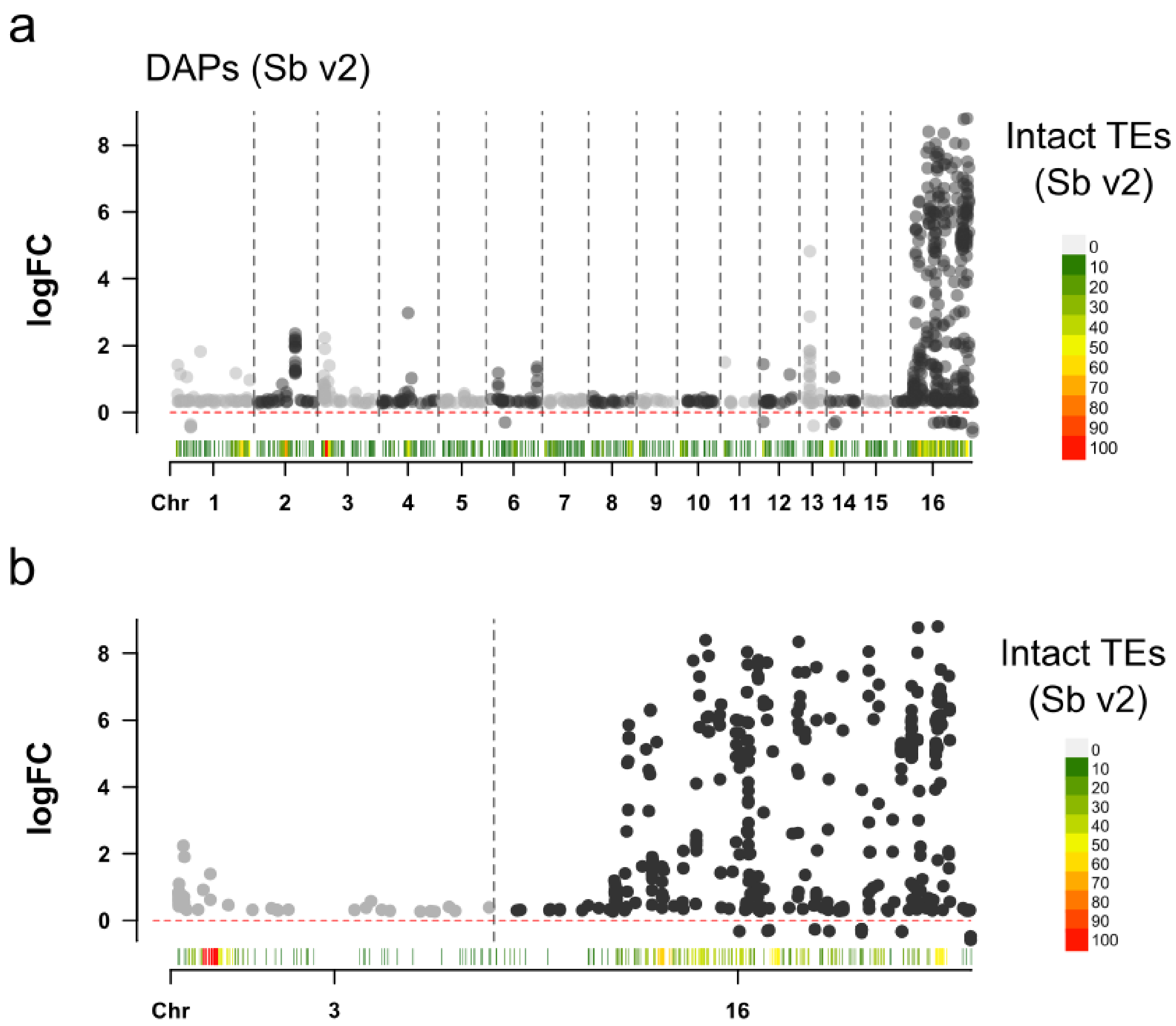


**Figure S2. Differential regions of accessible chromatin by supergene genotype are frequent near hotspots of intact TEs.**

**(a)** Peaks of differential chromatin accessibility based on alignment to long-read *Sb* v2 assembly. logFC values (positive indicating higher accessibility in *SB/Sb* relative to *SB/SB*) are shown for all differentially accessible peaks (DAPs) in worker brains. Color scales along the x-axis indicate the number of intact TEs (annotated with EDTA (Ou et al. 2019), see Methods) in 1e6 nucleotide bins across the genome. High numbers of DAPs are observed on chromosome 16 (Chr16) but and also near regions with dense TEs, including on Chr2 and Chr3. **(b)** Same as in (a) but for only Chr3 and Chr16 to highlight regions mentioned in the main text.

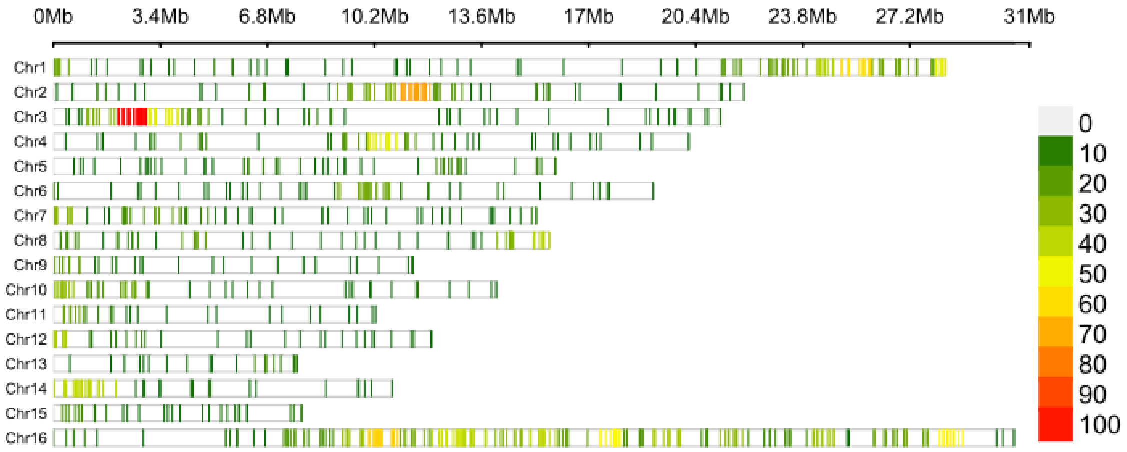


**Figure S3. Long-read genome assembly of *Sb* (v2) contains regions with high TE density.**

Colored bars across each chromosome indicate the number of intact TEs (annotated with EDTA (Ou et al. 2019), see Methods) in each 1e6 nucleotide bin across the genome of *Sb* v2 (a long-read based assembly). Particularly high numbers of TEs are observed in regions of Chr2 and Chr3, as well as throughout much of Chr16.

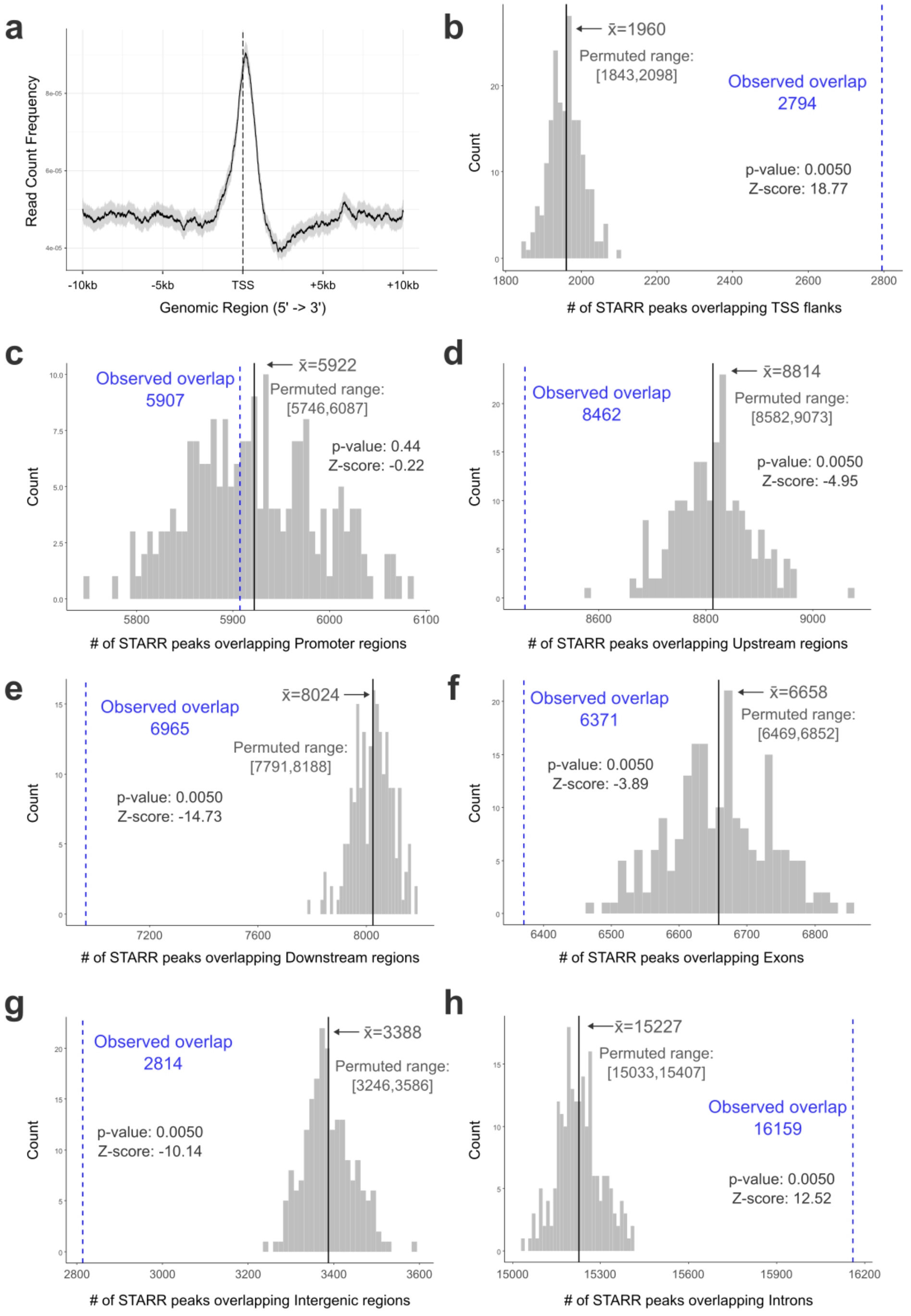


**Figure S4. STARR-seq peaks are frequently located near TSSs or within introns.**

**(a)** Frequency distribution of STARR peaks within 10kb of all annotated TSSs. Permuted overlap values (gray histogram) and empirical values (blue lines) for feature overlap of ATAC peaks with **(b)** TSS flanks (TSS +/- 200bp), **(c)** Promoters (0-5kb upstream of gene), **(d)** Upstream regions (0-10kb upstream of gene), **(e)** Downstream regions (0-10kb downstream of gene), **(f)** Exons, **(g)** Intergenic regions, and **(h)** Introns.

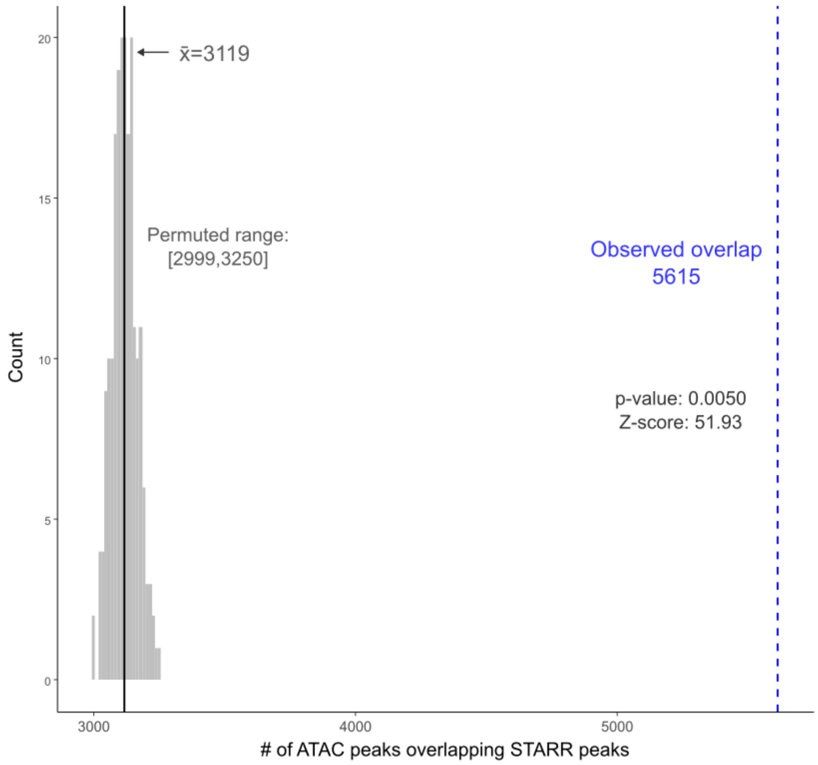


**Figure S5. ATAC-seq peaks are more likely to overlap STARR-peaks than expected by chance.**

Permuted overlap values (gray histogram) and empirical value (blue line) for overlap of ATAC-seq peaks with STARR-seq peaks, with n=200 permutations.

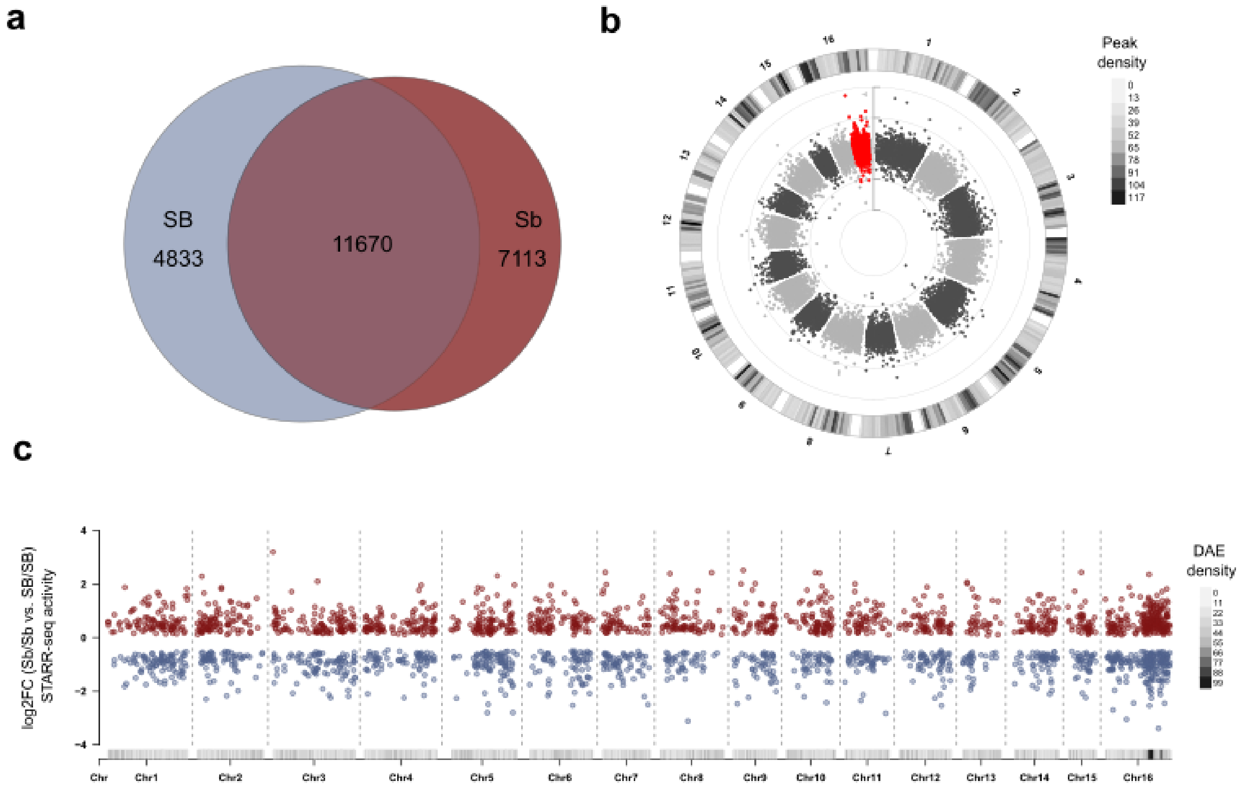


**Figure S6. STARR-seq identified enhancers were often shared across genotypes and located genome-wide. (a)** Of 23,616 active enhancers, 11,670 were shared between *SB* and *Sb* DNA fragments. **(b)** Enhancers were located genome-wide, including in the region of the supergene inversion (red dots). **(c)** Differentially active enhancers (DAEs) containing fixed allelic differences between *SB* and *Sb* fragments were located throughout the genome but with increased frequency near the inversion on chromosome 16, but were nearly evenly split between *SB*-biased and *Sb*-biased activity (~44% *SB*-biased and ~*56*% Sb-biased, Table S13).

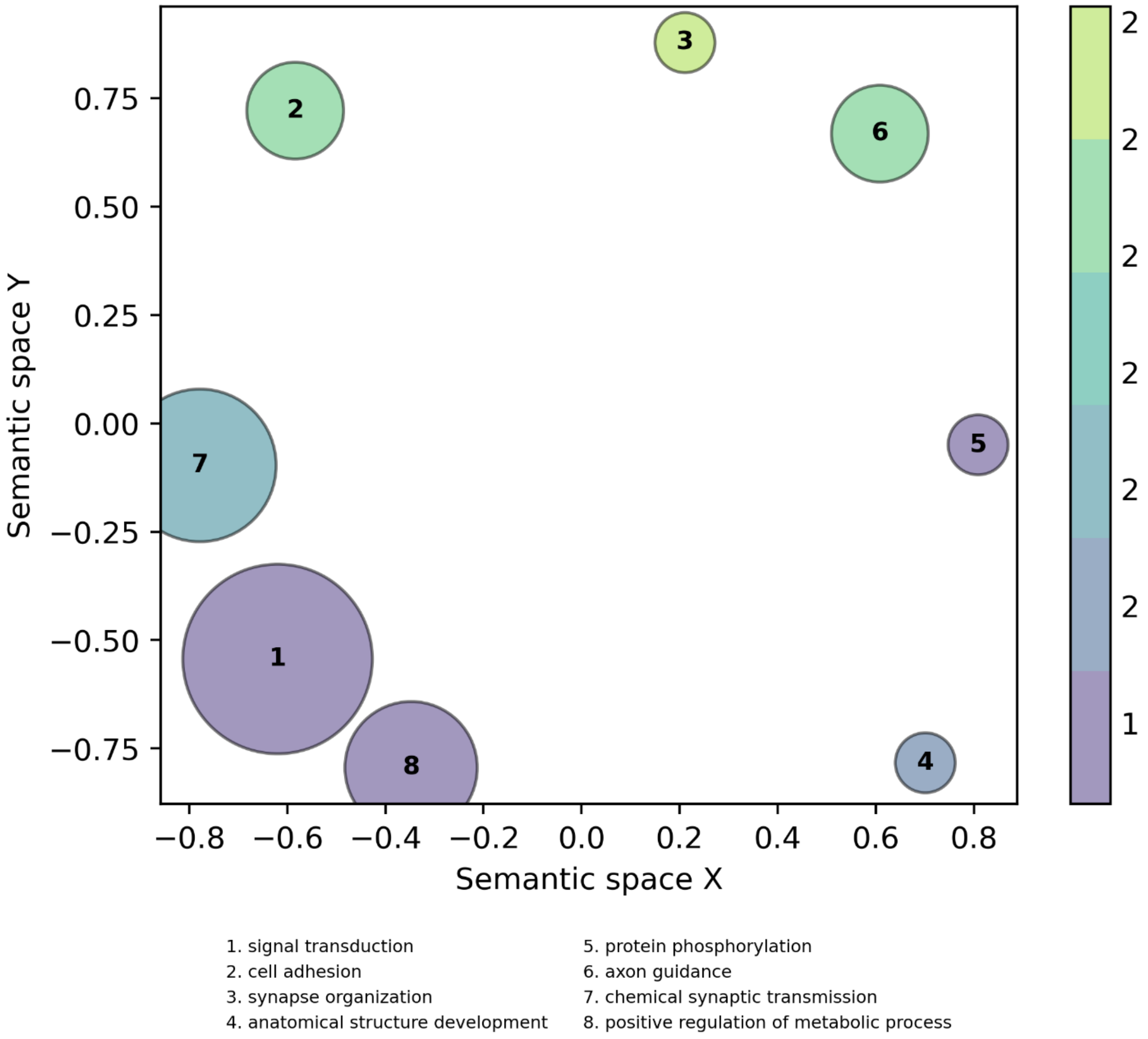


**Figure S7. Genes proximal to *SB*-biased DAEs were enriched for GO terms related to signal transduction, cell adhesion, synapse organization, and metabolism.**

Input genes included priority-assigned genes within 10kb of *SB*-biased DAEs and background set was all genes within 10kb of any enhancer. All significant (Benjamini-Hochberg corrected $p<0.05$) terms were clustered by semantic similarity using GO-Figure! (Reijnders & Waterhouse, 2021). Size of each circle is proportional to the specificity of the GO term (more specific = larger), and color indicates enrichment of each GO term in the input set relative to the proportion of genes in the background set.

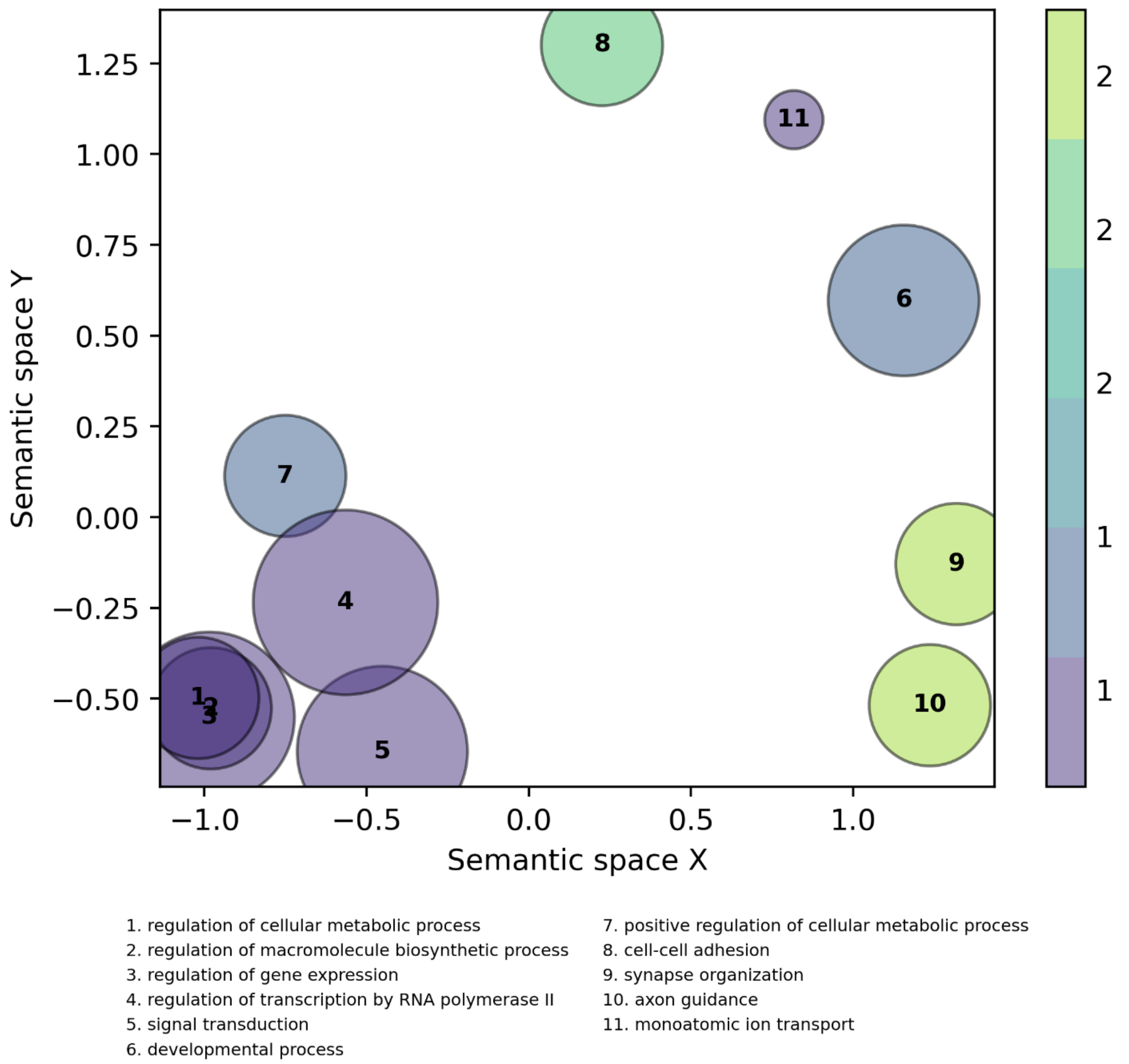


**Figure S8. Genes proximal to *Sb*-biased DAEs were enriched for GO terms related to metabolism, gene expression, cell-cell adhesion, and synapse organization.**

Input genes included priority-assigned genes within 10kb of *Sb*-biased DAEs and background set was all genes within 10kb of any enhancer. All significant (Benjamini-Hochberg corrected $p<0.05$) terms were clustered by semantic similarity using GO-Figure! (Reijnders & Waterhouse, 2021). Size of each circle is proportional to the specificity of the GO term (more specific = larger), and color indicates enrichment of each GO term in the input set relative to the proportion of genes in the background set.

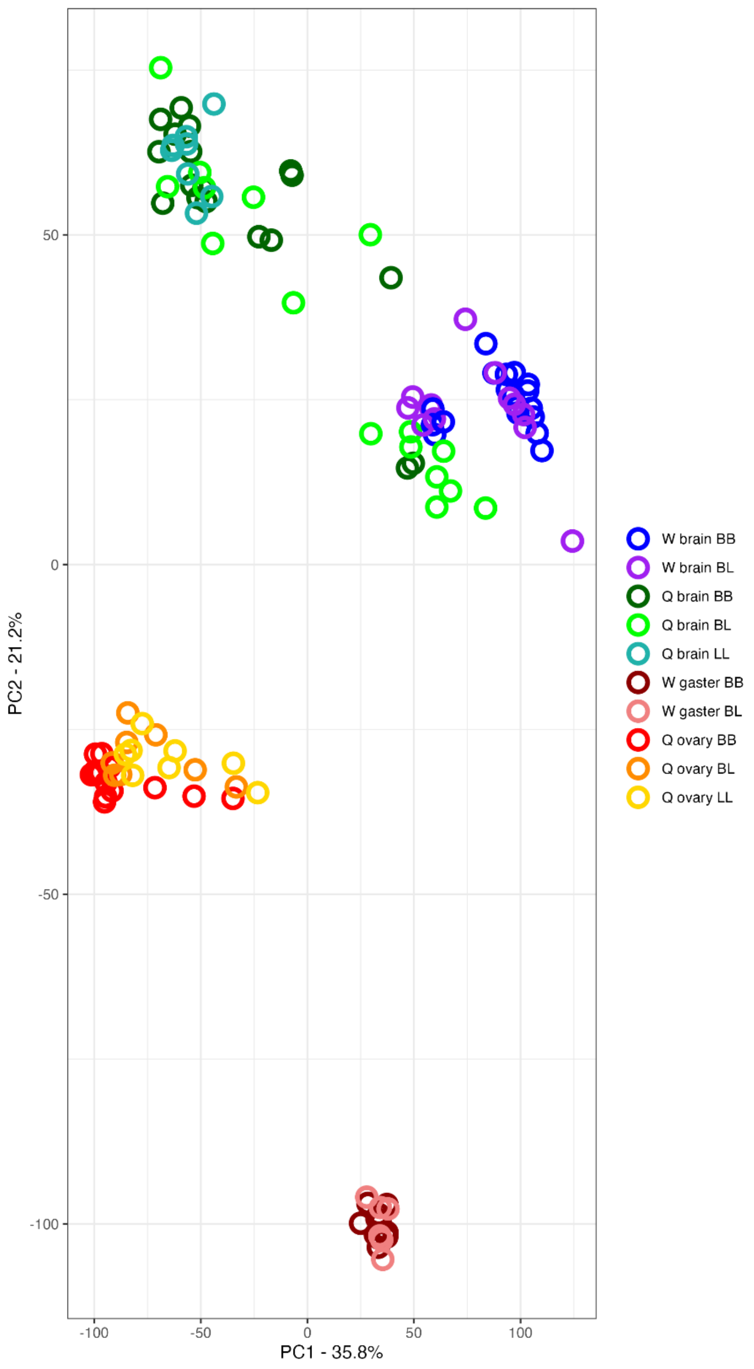


**Figure S9. Principal Component Analysis of RNAseq samples shows separation primarily by tissue type and caste, with limited separation by genotype.**

PCA of all RNA-seq sample used, which includes previously published individual bulk cell worker brain, worker gaster, queen (gyne) brain, and queen (ovary) samples (n=6-8 each) across *SB* (B) and *Sb* (L) genotypes. PC2 is most representative of tissue type, explaining 21.2% of variance in gene expression. No PC clearly representing genotype was identified.

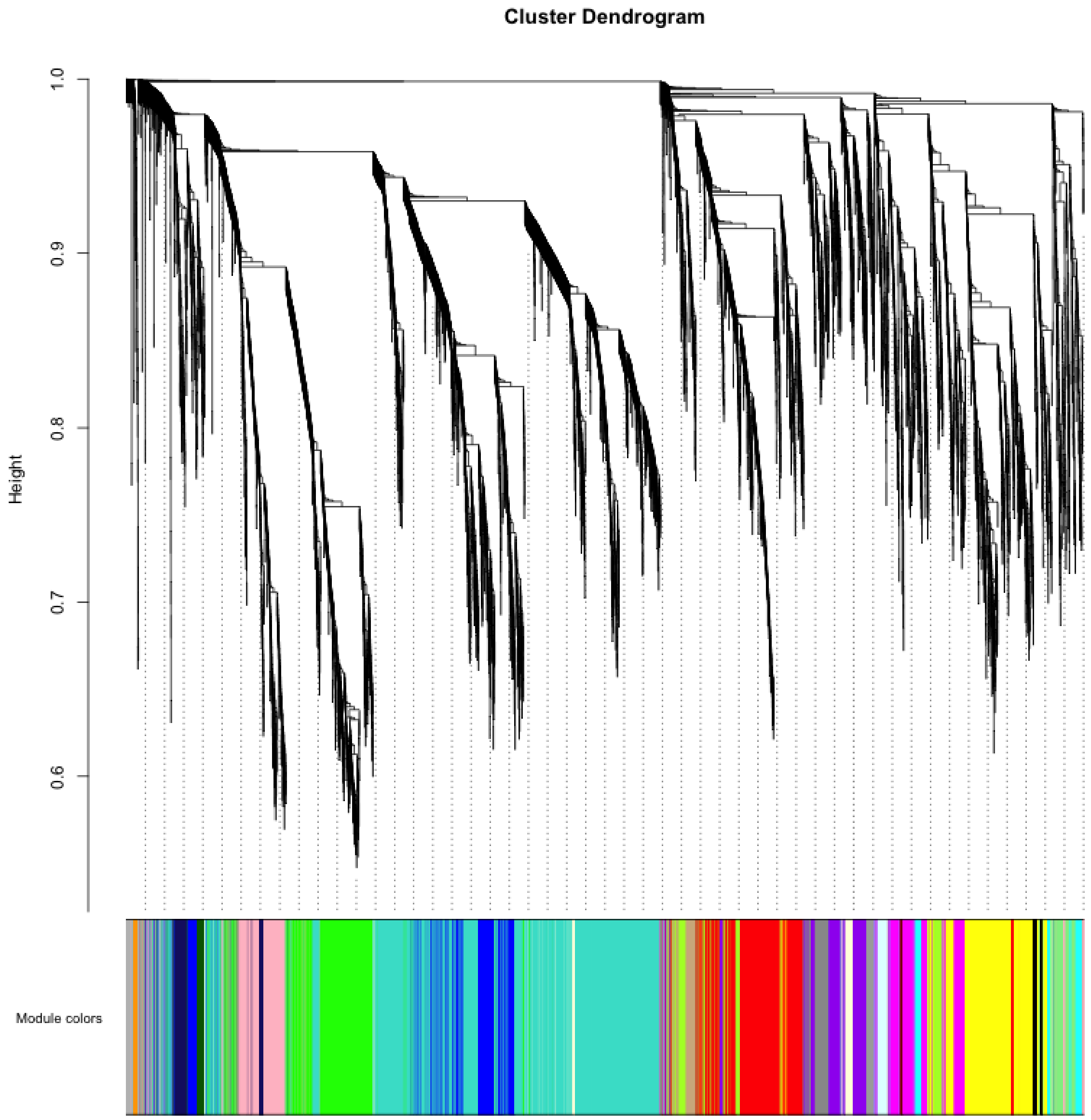


**Figure S10. WGCNA analysis identified 26 consensus modules.**

Results of agglomerative hierarchical clustering to identify WGCNA modules. Branches were cut using the hybrid dynamic tree cutting function with module detection sensitivity set to 2, minimum module size 20, and the cut height for module merging set to 0.15. Modules whose eigengenes were correlated above 0.85 were merged.

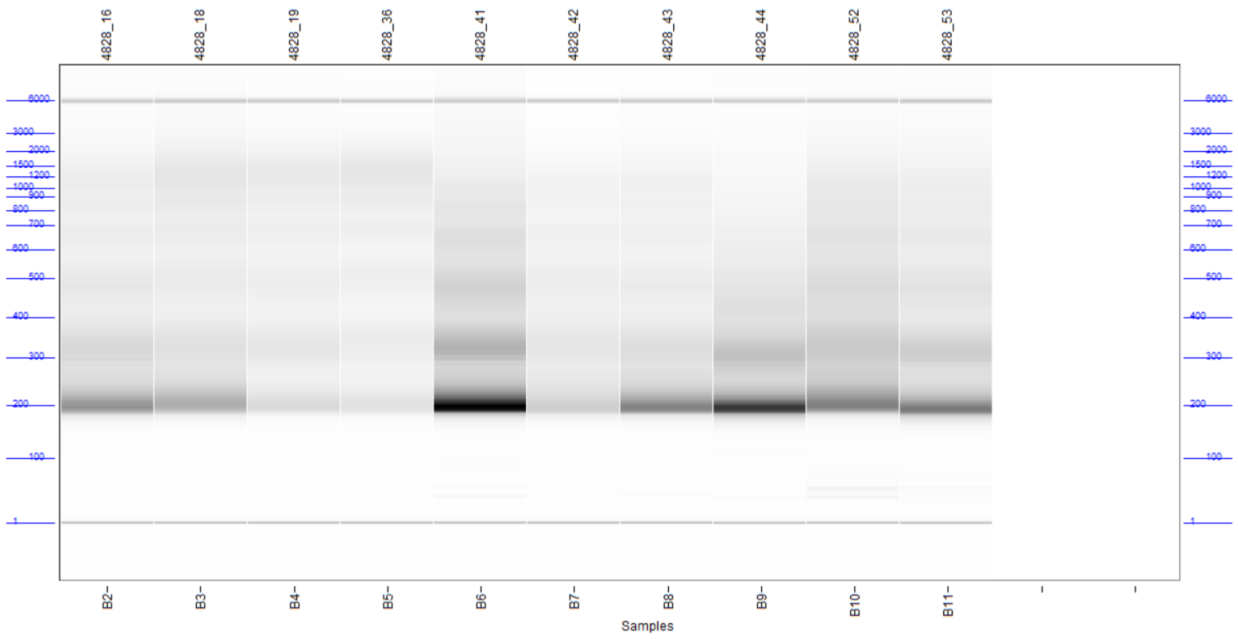


**Figure S11. Fragment analysis of ATAC-seq libraries demonstrates banding patterns corresponding with expected nucleosomal banding.**

ATAC-seq libraries selected for sequencing were screened for their nucleosomal banding patterns, corresponding to fragments containing 1, 2, 3, or additional nucleosomes with the ~147bp of associated DNA.

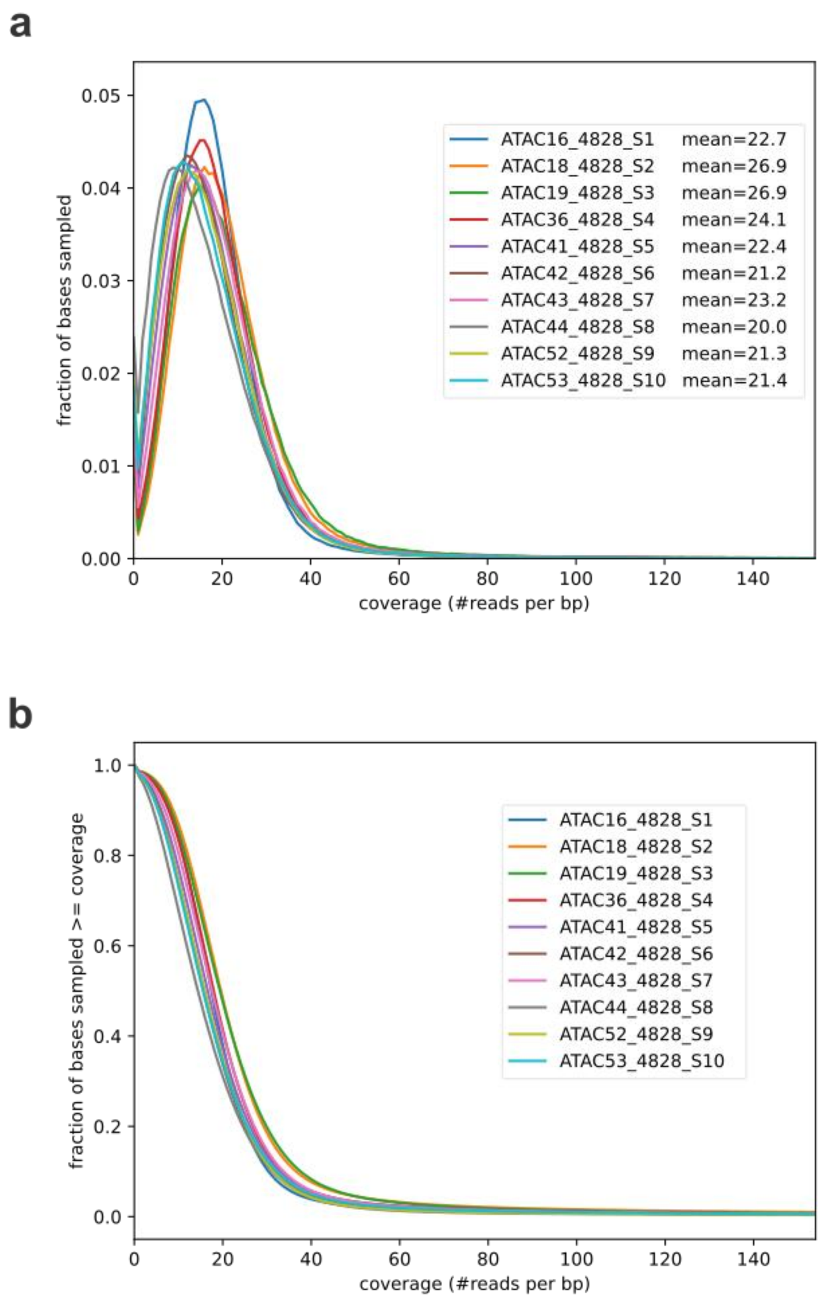


**Figure S12. ATAC-seq libraries had consistently high genomic coverage.**

**(a)** Fraction of bases in genome with denoted read coverage per sample. **(b)** Inverse cumulative frequency plot of genome coverage across ATAC-seq samples.

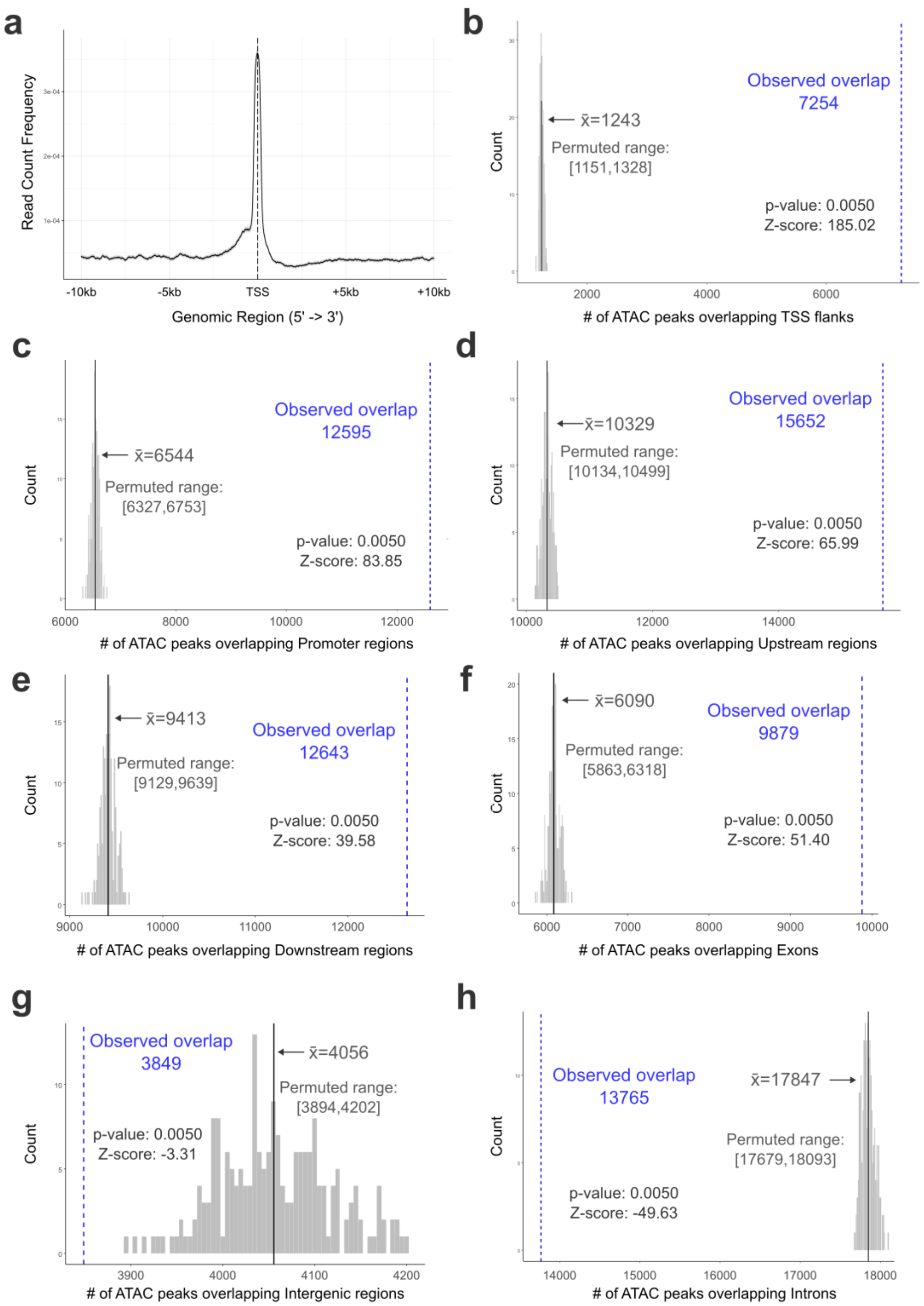


**Figure S13. ATAC-seq peaks are frequently located near TSSs and upstream of genes.**

**(a)** Frequency distribution of ATAC peaks within 10kb of all annotated TSSs. Permuted overlap values (gray histogram) and empirical values (blue lines) for feature overlap of ATAC-seq peaks with **(b)** TSS flanks (TSS +/- 200bp), **(c)** Promoters (0-5kb upstream of gene), **(d)** Upstream regions (0-10kb upstream of gene), **(e)** Downstream regions (0-10kb downstream of gene), **(f)** Exons, **(g)** Intergenic regions, and **(h)** Introns.

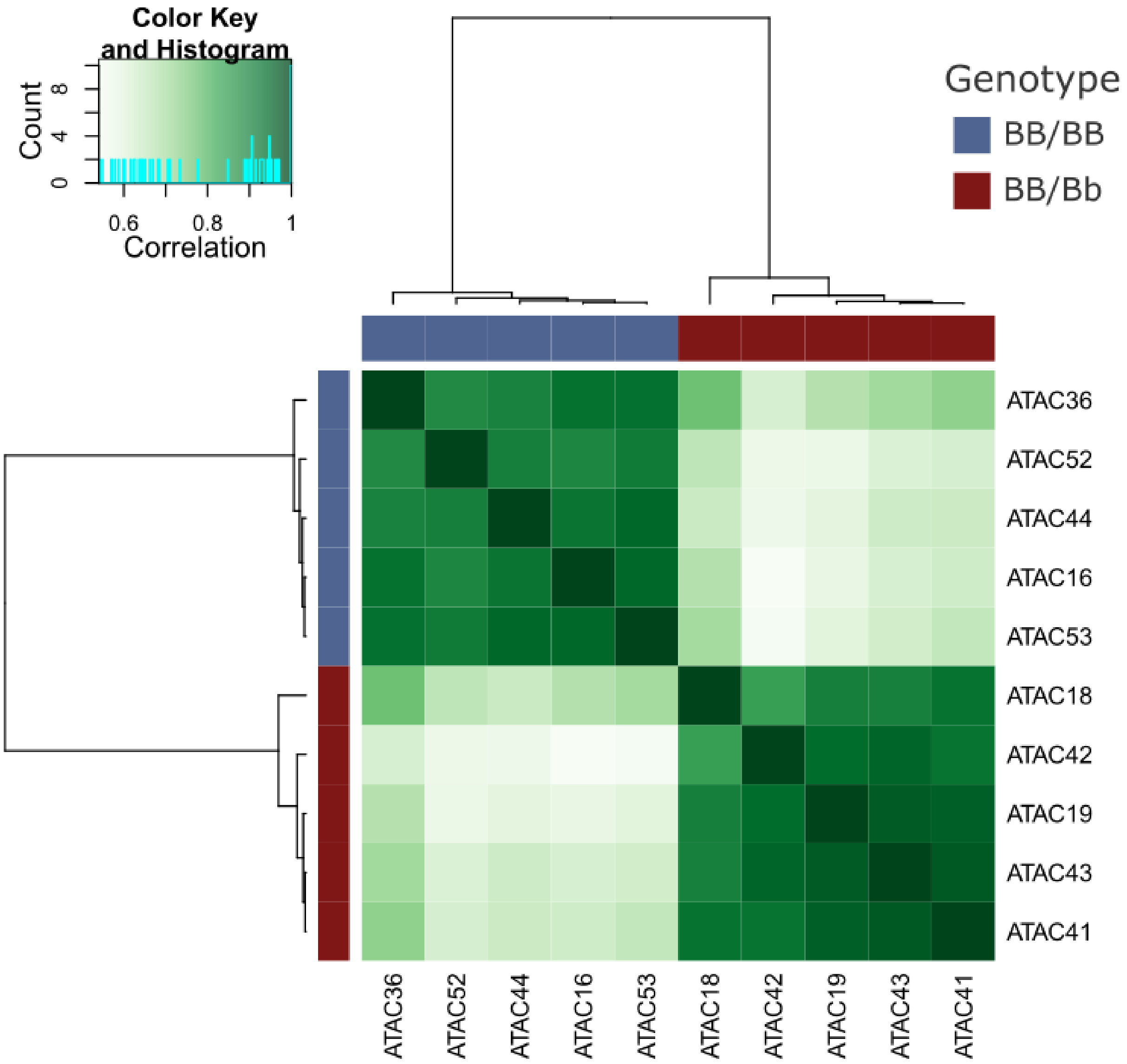


**Figure S14. Chromatin accessibility of DAPs strongly differentiates *BB/BB* and *BB/Bb* worker brains.**

Pearson correlation of differentially bound sites shows clear differentiation between genotypes. Plot produced with the dba.plotHeatmap function of DiffBind v3.12.0 (https://bioconductor.org/packages/release/bioc/html/DiffBind.html).
